# Bats decouple sonar gaze from steering to resolve sensory conflict

**DOI:** 10.64898/2026.08.08.743555

**Authors:** Nikita M. Finger, Shivam S. Chitnis, Grace Capshaw, Alara Kaplanoglu, Anand Krishnan, Cynthia F. Moss

## Abstract

When sensory modalities yield conflicting information, animals must rapidly reassess stimuli to select their actions. We induced auditory–visual conflict in free-flying echolocating Egyptian fruit bats, by fitting animals with prisms that shifted the perceived visual location of a landing perch while echoes returned from its veridical location. Bats that course-corrected within a single goal-directed flight did so by decoupling sonar gaze from steering, to enable rapid reweighting of visual and auditory cues. We designed artificial agents that used Bayesian inference to construct estimates of goal locations in their environment. When competing estimates directed active-sensing behaviors distinctly from steering, agents course-corrected more rapidly. Consistent with this idea, when bats were fit with prisms and earplugs that attenuated auditory localization cues, they were unable to course-correct. Removing prisms produced no systematic after-effects. Our framework suggests that instead of correcting their behavior after failure, animals could efficiently employ active sensing to resolve sensory conflict before failure occurs.

**Cover figure.**
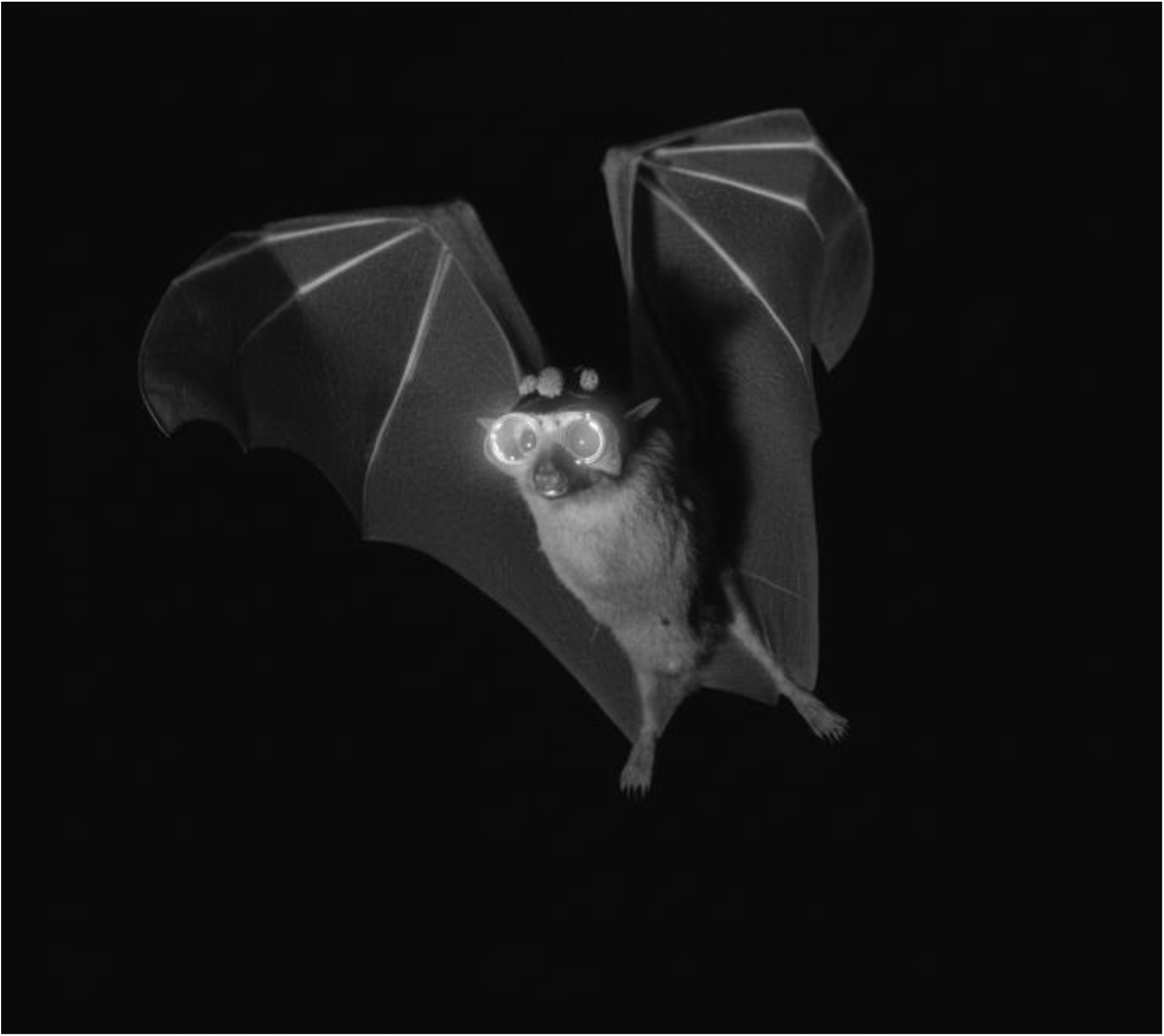
Bat wearing helmet with clear-glasses and tracking markers. Photograph © 2026 Nikita M. Finger / Moss Laboratory.

## Introduction

Animals rarely rely on a single sensory modality to guide action. Instead, they integrate information from multiple sensory cues (*1–3*) the relative weights of which can shift rapidly with environmental conditions, internal state, behavioral context, and task-specific cognitive demands (*4–8*). For instance, when presented with conflicting sensory information, animals often weight cues by their reliability (*9–11*). In visually-dominant animals, prism lenses have been employed to shift visual images relative to other sensory cues, revealing immediate visual capture by the shifted images, followed by gradual sensory and motor recalibration (*12–18*). This has also been demonstrated in locomotor tasks (*19*), where course correction is thought to reflect visual feedback control (*20*), rather than moment-to-moment changes in cue weights. However, in most species, the behavioral influence of distinct sensory cues is difficult to track during the course of navigation, leaving moment-to-moment cue reweighting largely unexplored.

We address this gap by leveraging the dual distal sensory systems of the Egyptian fruit bat, *Rousettus aegyptiacu*s, a highly visual animal that also uses tongue-click echolocation (*21–23*), to resolve rapid changes in sensory conflict during goal-directed flight. Egyptian fruit bats have large eyes, and show relatively high spatial resolution, low light-detection thresholds, and some binocular overlap, which supports visually-guided navigation (*24–26*). They also emit lingual sonar clicks, produced in pairs, directed off-axis to maximize auditory localization accuracy (*21, 22, 27*). The directional aim of biosonar serves as a measurable proxy for moment-to-moment spatial attention and information-seeking (*28–30*). For echolocating bats engaged in goal-directed flight, a stationary landing perch provides both visual and auditory spatial cues. These cues can be experimentally dissociated by fitting animals with prism lenses that shift visual images, while echolocation yields reliable auditory cues about the location of objects in the environment. The relative contributions of the two sensory modalities to ongoing behavior can be quantified by tracking the bat’s flight trajectory, head direction, and sonar beam aim as navigation unfolds.

During navigation, sensory information about the environment changes with self-motion, necessitating continuous updates to internal estimates of object and goal locations (*31–33*). Animals, as well as artificial agents, can control the spatial and temporal structure of sensory sampling, offering a powerful tool for directing how and when internal estimates are updated (*34–37*). Echolocating bats, for instance, regulate the rate and directional aim of their sonar emissions, enabling active sampling of specific regions of space (*24, 28, 38*). In doing so, they can validate or discount spatial estimates derived from conflicting sensory cues. By tracking the distinct sensorimotor outputs of bats confronted with conflicting spatial estimates of a target, we asked whether active sampling enables rapid cue reweighting to maintain goal-directed behavior.

We employed prism lenses to create auditory-visual conflict in Egyptian fruit bats that were trained to fly to a stationary landing target. Bats were tested with clear, dark, or prism lenses that preserved, occluded, or laterally displaced the apparent target location. In separate trials, bilateral plugs were inserted in the bat’s ear canals to attenuate echo feedback and probe the contribution of auditory processing to target localization when visual information was intact or altered. We hypothesized that bats use echolocation to resolve auditory-visual conflict during flight, initially by orienting toward the visually displaced target, before making a course-correction to the true target location when acoustic feedback is available. In parallel, we designed artificial agents within a Bayesian framework to test how active sensing could support this correction. We asked whether agents performed better when sensory sampling could be directed toward an alternative goal estimate instead of being coupled to the location currently guiding locomotion.

## Results

### Prisms reveal visual displacement perception and rapid course-correction during flight

We tested whether prism lenses biased head orientation and steering in flight (Fig. 1A). Bats wearing prism lenses initially steered and oriented in the direction of the 23° visual displacement: left with left-shifting prisms and right with right-shifting prisms (χ^2^(6) = 34.54, p < 0.001; χ^2^(54) = 133.88, p < 0.001; Fig. 1B-J, Fig. S1). The smaller magnitude (11°) right-shifting prism lenses produced a similar but weaker rightward trajectory and head orientation bias (Fig. S1). Over the course of each trial, bats directed their flight paths and head direction back toward the veridical stationary target, course-correcting successfully even on the first trial of a session (χ^2^(18) = 126.60, p < 0.001; χ^2^(42) = 70.01, p = 0.004; Fig. 1E-J, Fig. S2). As a result, they often landed on the target when wearing prisms (Fig. S3A). Larger prism shifts did, however, reduce direct landings by ∼25%, increase back-and-forth landings by ∼5-10% (χ^2^(18) = 243.95, p < 0.01, Cramer’s V = 0.339; all p_Holm_ < 0.01, Fig. S3A), and, as with smaller shifts, decrease the straightness of bats’ flight trajectories (Fig. S3B).

**Figure 1.**
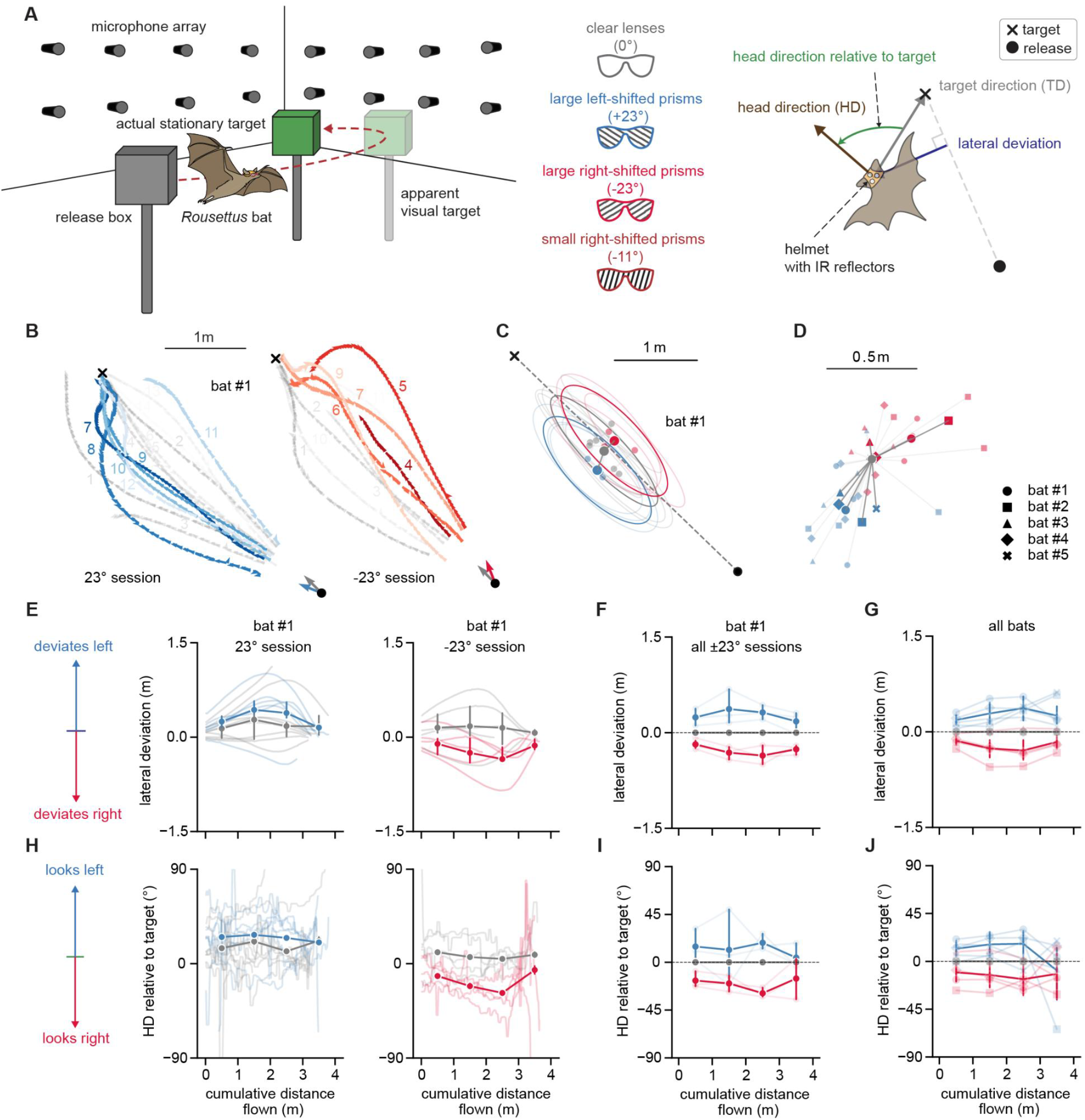
Auditory-visual conflict assay in freely flying Egyptian fruit bats. A) Schematic of free-flight behavioral task, experimental setup, lens conditions, and variables extracted for analysis. B) Representative flight trajectories from bat #1 during a representative left (blue) and right (red) prism session. Gray trajectories represent clear-lens (control) trials. Numbers indicate trial order. C) Centroids of flight trajectories for bat #1 under left and right prism conditions, showing the average lateral displacement of flight paths relative to control. Ellipses indicate spread across trials. D) Mean centroids relative to control for all bats under left and right prism conditions, illustrating consistent prism-dependent lateral shifts across individuals. E–G) Lateral flight deviation across cumulative distance flown, showing mean and 95% CI. E) Representative single-session data from bat #1 illustrating low within-session variability. F) Pooled data for bat #1 across all sessions normalized to clear-lens trials, illustrating low across-session variability. G) Pooled data across all bats normalized to clear-lens trials. H–J) Similarly, for head direction relative to the target across cumulative distance flown.

### Bats increase biosonar sampling when vision is displaced or unavailable

Bats strongly modulated their sonar click rate by adjusting the interval between consecutive click pairs (inter-pair interval) across lens conditions (χ^2^(54) = 98.69, p < 0.001; Fig. S3C–D). A small 11° visual shift produced only a modest, non-significant increase in sonar click rate (p_Holm_ = 0.506). Larger 23° shifts elicited higher click rates, reaching significance for the large right-shifting prism (p_Holm_ = 0.017) and showing the same directional trend for the large left-shifting prism (p_Holm_ = 0.146). Bats clicked the most when vision was occluded during dark-lens trials (all p_Holm_ < 0.001, Fig. S3D). Bats tended to fly slower when visual information was disrupted, with the greatest contrast observed between the dark- and clear-lens conditions (Fig. S3B). Despite this trend, lens condition did not significantly influence flight speed (χ^2^(6) = 1.92, p = 0.927). As bats approached the target, they reduced their flight speed (χ^2^(3) = 137.52, p < 0.001, Fig. S3B), and increased sonar click rate (χ^2^(49) = 315.32, p < 0.001, Fig. S3D), but did not change sonar behavior systematically across repeated trials within a day (χ^2^(7) = 3.59, p = 0.826, Fig. S3E). This trial-related pattern did not differ across conditions (χ^2^(6) = 0.56, p = 0.997, Fig. S3E).

### Head direction and sonar aim diverge during sensory conflict trials

Sonar direction (Fig. 2A) relative to target direction (SD-TD) and head direction (Fig. 1A) relative to sonar direction (HD-SD) changed over the course of each trial (HD-SD: χ^2^(54) = 210.20, p < 0.001; χ^2^(42) = 168.35, p < 0.001; SD-TD: χ^2^(54) = 173.58, p < 0.001; χ^2^(42) = 116.56, p < 0.001, Fig. 2B). Early in each trial, bats aimed their sonar closer to the true target location, whereas their head direction followed the prism-shifted visual target. Later in each trial, bats shifted the sonar aim toward the visually displaced target, while reorienting head direction toward the true target location (Fig. 2A-C). This reversal reflected dynamic changes in the click-pair pattern: instead of the typical left-right alternating click pattern observed in clear-lens trials, bats produced same-side click pairs which were directed left or right as they approached the target. For example, in left-shifted prism trials bats initially produced right-right click-pairs, and then later in the trial they produced left-left click-pairs. The opposite pattern was observed for the right shifted trials (Fig. 2A). Sonar-head divergence (HD-SD) far from the target (> 1.5m and < 2.5m), was inversely correlated with the maximum lateral deviation of the bat’s trajectory, suggesting that aiming the sonar at the actual target at the beginning of the trial facilitated course correction (Fig. 2F). Sonar-head divergence and its correlation with maximum deviation were not prominent in trials without prisms (Fig. S4), suggesting that the bat decoupled sonar aim and heading primarily upon detecting conflict.

**Figure 2.**
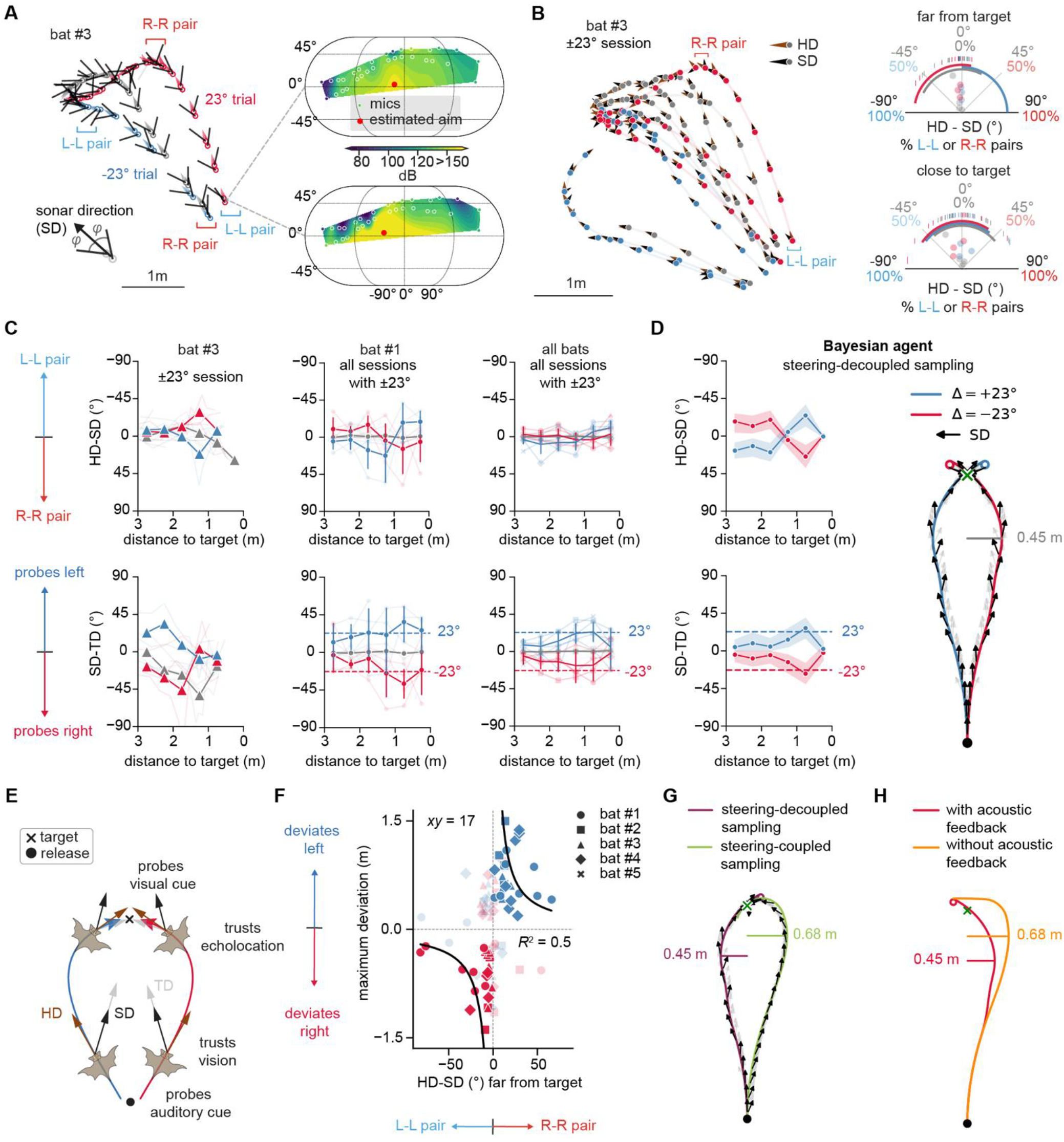
Biosonar aim diverges from head direction during audio-visual conflict. A) Extracted sonar direction of each click with representative power spectral density (PSD) overlaid on a representative left-shifted and right-shifted prism trial from a single session. Note the presence of “left-left” (L-L) and “right-right” (R-R) clicks-pairs. B) Representative prism-shifted trajectories showing head direction (HD) and sonar direction (SD; left) and corresponding HD-SD and proportion of L-L vs. R-R click-pairs as a function of distance from the target (right) for a single session with both left and right prism trials.

Ticks represent indidividual click-pairs and circles represent individual trials. C) HD-SD (top) and SD-TD (bottom) across distance from target, showing mean and 95% CI for data from a representative session from bat #3, pooled data for bat #1 across all sessions normalized to clear-lens trials, and pooled data across all bats normalized to clear-lens trials. D) HD-SD, SD-TD and trajectories with sonar direction for the model Bayesian agent showing every 3^rd^ sonar pair in black. Note how sonar deviates from heading. E) Summary schematic illustrating how HD, SD, and TD evolve over a single flight. F) Control-normalized maximum deviation within a trial vs. HD-SD far from the target. Dark points indicate trials that match the expected trend (HD-SD < 0° and rightward deviation for the right prism and HD-SD > 0° and leftward deviation for the left prism). Curve fit to the dark points shows an inverse relationship between early sonar-head divergence and maximum lateral deviation. G) Model trajectories with and without coupling of sampling to steering. Naive strategy in which sampling is aligned with steering leads to larger deviation. H) Model trajectory with and without acoustic feedback. Without feedback, the agent fails to approach the target.

### Decoupling information-seeking from steering allows for rapid course correction

To explore the potential benefits of the bat’s adaptive behaviors, we modeled a Bayesian agent that maintained a posterior belief over target locations in a world-centered coordinate frame and used it to execute low-level policies (*32*, Fig. 2D). The agent constructed a working hypothesis about the target’s location by computing the most likely location of the target (Fig. S7, S8), which guided steering. A sampling policy that evaluated prominent, but less likely, locations of the target reproduced observed sonar-head divergence behavior (Fig. 2D-E). Switching to the naive strategy of probing the most-likely hypothesis increased lateral trajectory deviation (Fig. 2G). Sampling the veridical target at the beginning of the trial rapidly modified the posterior, facilitating timely course correction (SI Movie 1). A utility function that penalized mis-localizing target estimates offered a decision-theoretic explanation for probing alternate hypotheses (SI Movie 2). Further, eliminating acoustic feedback prevented corrective steering, suggesting that plugging the bats’ ears might prevent them from approaching the target (Fig. 2H, SI Movie 3).

### Bats wearing prisms could not course-correct with reduced acoustic feedback

Bats fit with earplugs that significantly attenuated their hearing sensitivity failed to correct their prism-shifted trajectory (prisms +EP° vs. clear: p_Holm_ < 0.01, Fig. 3A-D, F). This failure was not attributable to perturbed visual input: when acoustic feedback was available during prism trials, bats were able to approach and land on the target (χ^2^(6) = 66.763, p < 0.01; prism vs. clear: p_Holm_ < 0.01, Fig. S2, S3A). They landed even in dark-lens conditions where visual input was removed, flying along straighter trajectories than under clear-lens conditions (Fig. 3A-D, Fig. S5A). In contrast, when bats were fit with both dark lenses and earplugs that attenuated echo feedback, they failed to land on the target (Fig. S5A), showing that they indeed used echolocation rather than spatial memory to course-correct in the prism trials. Neither was the failure caused by reduced acoustic feedback alone: bats fit with earplugs and clear lenses landed directly on the target at rates similar to clear-lens controls (EP vs. clear: p_Holm_ = 0.165; landing type: χ^2^(3) = 3.37, p_Holm_ = 0.675, Fig. 3A-D, Fig. S5A). Consistent with this finding, attenuating auditory feedback itself did not alter click rate (p_Holm_ = 1.00), whereas bats fit with both prisms and earplugs tended to increase click rate relative to clear-lens controls (p_Holm_ = 0.076, Fig. 3E).

**Figure 3.**
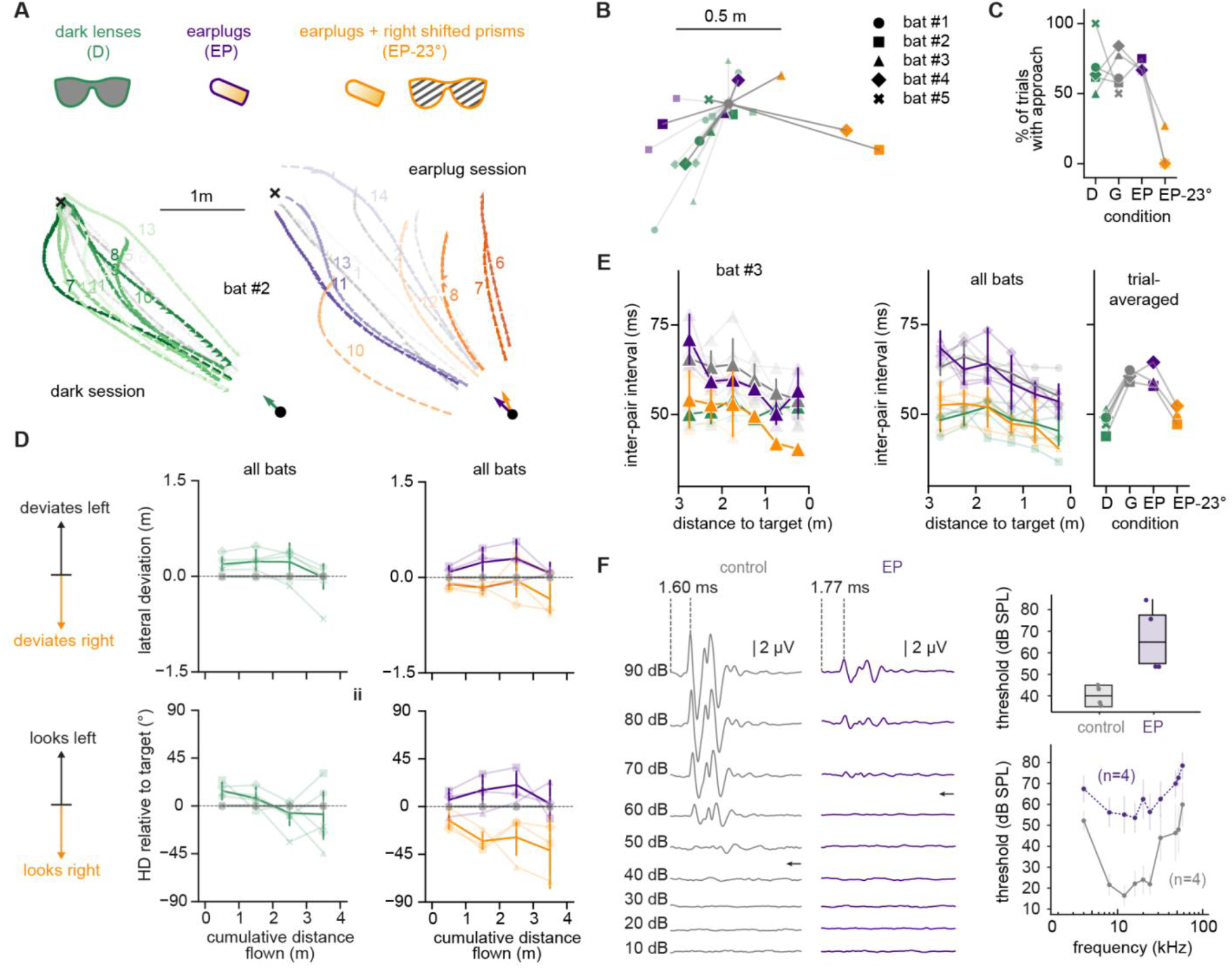
Attenuated acoustic feedback dramatically impairs navigation when bats view the world through prisms. A) Experimental lens and earplug conditions used to test the contribution of echo feedback to navigation and representative flight trajectories from a single bat under dark-lens (green), earplug-only (purple), and earplugs+prism (orange) conditions. Gray trajectories show clear-lens control trials from the same day. X marks the location of the stationary target. Numbers indicate trial order. B) Mean centroids relative to control, illustrating condition-dependent shifts in flight direction across individuals for each session. C) Percent landing success for each bat under the three experimental conditions (colored points) and control (gray points) conditions, demonstrating the strongest reduction in success occurred when visual conflict was combined with reduced acoustic feedback. D) Lateral deviation (top) and head direction relative to the target (bottom) across cumulative distance flown across experimental conditions for all bats, normalized to clear-lens trials. Positive values indicate leftward deviation; negative values indicate rightward deviation. E) (Left) Inter-pair interval as a function of distance to target recorded from a single bat, mean inter-pair interval as a function of target distance across all bats and (right) trial-averaged inter-pair interval across conditions, illustrating changes in ICI under degraded sensory conditions. F) Sound-evoked auditory brainstem response (ABR) recordings show diminished hearing sensitivity with earplugs (EP) relative to control measurements, including reduced and delayed ABR waves (waveforms evoked by broadband clicks from 10-90 dB SPL; threshold indicated by arrow), and elevated thresholds to clicks (top boxplot) and tones (bottom audiogram).

### Experience improves course correction over successive trials

We next tested whether bats adjusted their navigation with repeated prism exposure across trials. The magnitude of lateral deviation was different between early-and late-prism trials (Fig. 4A-B, D). The rate of learning differed between prism displacement conditions, with bats requiring fewer trials to learn to execute control-like trajectories for the 11° right-shifting prism than for the 23° left or right prisms (Fig. 4C). Landing success also improved over trials (χ^2^(1) = 5.721, p = 0.017). Head–sonar alignment tightened across successive flights, indicating that orienting and acoustic sampling became more coordinated with experience (χ^2^(6) = 14.90, p = 0.021, Fig. S6A). Sonar aim tended to become more target-directed over a session, although this effect did not differ significantly across lens conditions (trial effect: χ^2^(7) = 16.56, p = 0.020; condition × trial: χ^2^(6) = 9.01, p = 0.173). Specifically, sonar aim far from the target became more directed at the true target over the course of the session (Fig. S6B). Sonar aim close to the target remained directed at the apparent visual target over the course of the session (Fig. S6B) suggesting that the bat’s strategy remained unchanged.

**Figure 4.**
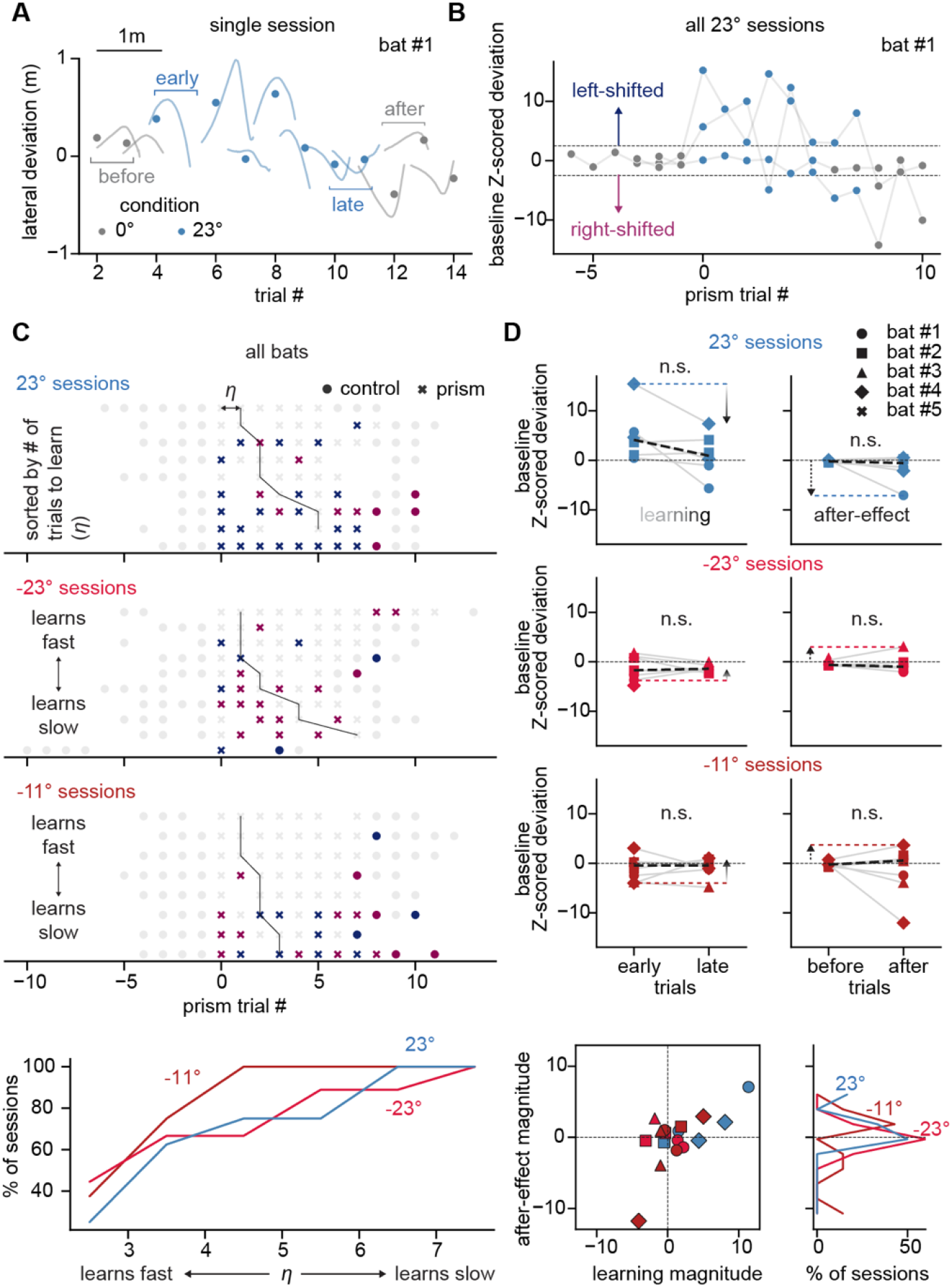
Trial-by-trial learning during prism exposure and the absence of after-effects. A) Example single-session trajectory deviations from bat #1 during a 23° prism session. Gray curves show clear-lens control trials before and after prism exposure; blue curves show prism trials. B) Trial-by-trial change in baseline-normalized deviation across all 23° sessions for bat #1. Values greater than 1.5 indicate left-shifted trials, less than 1.5 indicate right-shifted trials, and the rest are unshifted C) (Top) Session-by-session summary of the rate of learning across all bats for 23° left, 23° right, and 11° right prism sessions. Colored points show left-right- or un-shifted trajectories. (Bottom) Cumulative distributions of trials to learn for the three prism conditions. D) (Top) Session-level summaries of learning magnitude and after-effect. Left column: change from early to late prism trials (“learning”) for 23° right, 23° left, and 11° right sessions. Right column: change from before-prism to after-prism clear-lens trials (“after-effect”) for the same prism conditions. All comparisons were non-significant (n.s.; Wilcoxon signed-rank test, all p>0.05). Points represent individual sessions; dashed lines indicate the group means. (Bottom) Relationship between learning magnitude and after-effect magnitude across sessions and distribution of after-effect magnitudes for each prism condition, showing weaker within-day after-effects relative to learning expressed during prism exposure.

### Bats correct flight trajectories without exhibiting classical prism after-effects

If bats recalibrated their sensorimotor coordination with experience, removing prisms should have produced systematic opposite-direction after-effects in subsequent clear-lens trials (*19, 39*). We found no systematic evidence of such after-effects (Fig. 4D). Trajectory deviation (Fig. 4D), head direction, and sonar direction did not differ between clear-lens trials before- and after-prisms (trajectory: χ^2^(1) = 0.002, p = 0.969; head direction: χ^2^(1) = 1.661, p = 0.197; sonar direction: χ^2^(1) = 1.220, p = 0.269). These before-vs-after changes also did not depend on prism condition (trajectory phase × prism type: χ^2^(2) = 0.896, p = 0.639; head direction phase × prism type: χ^2^(2) = 0.796, p = 0.672; sonar direction phase × prism type: χ^2^(2) = 3.741, p = 0.154). When we compared the magnitude of the before-vs-after change in lateral deviation against the early-vs-late change, we uncovered a correlation (Pearson R = 0.65, p = 0.003, Fig. 4D), suggesting that an experimental paradigm with long-term prism exposure might reveal systematic after-effects by inducing recalibration.

## Discussion

### Rapid resolution of sensory conflict in flight

Free-flying Egyptian fruit bats resolved sensory conflict during goal-directed navigation within a single flight when prism-displaced visual images and veridical echoic target locations specified competing target positions. Early in each prism trial, flight trajectories and head direction shifted toward the direction of the visually displaced target, consistent with visual dominance in Egyptian fruit bats (*24*), whereas biosonar probed the veridical target. Later in each trial, trajectories and head direction reoriented toward the veridical target, as biosonar sampled the prism-shifted location.

### Simultaneous expression and evaluation of competing cues as navigation unfolds

To navigate to a stationary target when vision and echolocation cues conflicted, bats rapidly reweighted cues online, flying towards one sensory cue while actively probing another with biosonar. Rather than undergoing sensorimotor recalibration or collapsing conflicting stimuli into a fused percept, bats solved a causal-inference problem by maintaining competing hypotheses about conflicting cues (*11, 33, 40, 41*). Flight trajectories reflected the cue guiding locomotion, whereas the sonar aim reflected an information-seeking process directed toward its competitor, demonstrating that the cue being sampled was not always the cue driving locomotion. This strategy was effective in rapidly resolving sensory conflict, as sonar aim divergence from heading was inversely correlated with trajectory deviation. Decoupling steering and active sampling in a Bayesian model reduced trajectory deviations. By validating or discounting alternative hypotheses, bats reassign cue weighting as evidence accumulated. This behavioral dissociation parallels recent work in rodents, suggesting that the nervous system can internally sample locations distinct from the animal’s current heading or movement direction (*42, 43*). Together, these findings point to a broader principle: navigation may benefit from evaluating spatial locations without steering towards them (*44*).

### Experience refines information-seeking

Across repeated prism trials, landing success increased and head–sonar coordination tightened, suggesting that bats learned to sample information more effectively. This improvement was not explained by a general increase in sonar click rate, which did not change systematically across trials. Instead, experience shaped the priority assigned to competing cues: governing which cue guided navigation, but also which cue was sampled by sonar and hence the rate at which new evidence modified flight control.

### Active sampling is required for sensory conflict resolution

Bats modulated acoustic sampling when visual information was unavailable or in conflict with auditory information. Consistent with prior work showing increased biosonar use under low-light conditions (*24*), bats increased sonar click rate in darkness and with larger prism shifts. Active sampling was required for sensory conflict resolution: bats wearing prisms that shifted visual images and earplugs that attenuated auditory cues failed to reliably course-correct or land; whereas earplugs alone did not impair navigation. Therefore, auditory feedback was not required for navigation under natural conditions but was necessary to resolve sensory conflict.

### Online cue reweighting, not recalibration

The bats’ behavioral response to prism shifts of visual images differed from classical prism adaptation. Prism perturbations have long been used to study sensory or sensorimotor recalibration following visual capture, evidenced by the consistent after-effects that emerge after prism removal (*18, 39*). Prism exposure can also alter locomotor trajectories, generalize across movements, and recalibrate auditory spatial maps over longer timescales (*16, 17, 19, 45, 46*). In motor-learning frameworks, online feedback control updates ongoing movements, whereas repeated errors can recalibrate feedforward motor plans over longer timescales (*47–49*). Here, course correction unfolded as the bat approached the landing perch while the conflict persisted and produced no systematic within-day behavioral after-effects. Although longer prism exposure may uncover a slow-timescale recalibration, the behavior reported here is better explained by the ongoing evaluation of competing cues.

## Conclusion

Our findings offer novel advances to the weighting of sensory cues used to guide navigation. Visual and auditory cues were not combined to obtain a single spatial estimate guiding navigation, nor were they reweighted solely in response to landing failures. Instead, bats maintained competing estimates of the target location based on different sensory cues: one cue guided locomotion, the other information-seeking. This finding inspires future research to determine if this strategy generalizes across biological and artificial sensing systems in which agents must act when stimulus information conflicts or is ambiguous. Quantitative analysis of action selection and information seeking thus reframes sensory conflict from a problem of error correction after failure to one of online inference before failure occurs.

## Acknowledgments

James Garmon for designing the helmet and interchangeable lenses. Jake Sayles for help with troubleshooting equipment. Ann Hermundstad for useful discussions on Bayesian modeling. Christopher Honey for manuscript advice. Data Collection and Analysis: Wendy Chen, Magdiel Jaroszewski, Keegan Eveland, John Cole, Myles Gosha, Yiwen Lui, Laura Panlilio, DaQuan Greenwell, Mariana Meade, Jennifer He, Maxwell Rho, Irene Park, Jose Jarquin, Racheal Timothy, Leslie Bucio.

## Funding

The Center for Hearing and Balance T32 Postdoctoral Fellowship T32DC000023-39 (NMF).

National Institutes of Health R01 NS121413 (CFM)

Human Frontiers Science Program Research Grant RGP0045/2022 (CFM)

National Science Foundation Brain Initiative Grant NCS-FO 1734744

(CFM) National Science Foundation CRCNS Grant 2011619 (CFM)

Office of Naval Research Grants N00014-23-1-2086 and N00014-17-1-2736 (CFM)

## Author Contributions

Conceptualization: NMF, SSC, CFM, ANK, GC, ALK

Data curation: NMF, GC, SSC, ALK

Formal analysis: SSC, NMF, GC

Funding acquisition: CFM, NMF

Investigation: NMF, SSC, GC, ALK

Methodology: SSC, NMF, GC, ANK, ALK, CFM

Project administration: NMF, CFM, SSC

Resources: CFM

Software: SSC, NMF, GC

Supervision: CFM, ANK

Validation: SSC, NMF, GC

Visualization: SSC, NMF, GC

Writing – original draft: NMF, SSC, GC

Writing – review & editing: CFM, ANK, ALK

## Diversity, equity, ethics, and inclusion

NA

## Competing interests

Authors declare that they have no competing interests.

## Data, code, and materials availability

Data and analysis code will be made publicly available through a GitHub repository upon publication

## Notes

### Competing Interest Statement

The authors have declared no competing interest.

