## Supplementary Materials for "Bats decouple sonar gaze from steering to resolve sensory conflict"

**The PDF file includes:**

Materials and Methods
Supplementary Text — Theory (separate PDF)
Figs. S1 to S8
Tables S1

**Other Supplementary Materials for this manuscript include the following:**

Movies S1 to S3

#### Materials and Methods

Subjects

We tested five adult Egyptian fruit bats (*Rousettus aegyptiacus*, 2 females, 3 males). Animals were housed and handled under protocols approved by the Johns Hopkins University Institutional Animal Care and Use Committee (IACUC) and adhered to the National Institutes of Health guidelines (Guide for the Care and Use of Laboratory Animals, 2011), the U.S. Animal Welfare Act, and the Public Health Service Policy. The university maintains full AAALAC accreditation.

Behavioral Paradigm

We conducted experiments in a large (6m x 6.8m x 2.5m) acoustically attenuated flight room. (Fig. 1a). We trained bats to fly from a release box to a landing target while wearing a custom-made lightweight (11g, accounting for approx.≤10% of the bat's body weight) helmet with interchangeable lenses affixed using magnets. The landing target was a green, acoustically reflective box (30cm x 30cm x 30cm) that was centered on the far wall at a fixed height of 1.25 m.

We tested bats' performance in the flight task under five visual conditions (Fig. 1b): clear lenses (control), left-shifting prisms (23°, 3M™ Press-on Fresnel Prisms, 1mm thickness), right-shifting prisms (23°), smaller right-shifting prisms (11°), and dark (opaque) lenses. Each daily session comprised 13–16 trials, the first 2-5 trials of which were always performed with clear lenses to establish baseline performance for each bat. The middle 5-9 trials included the day's test condition in which bats performed the flight task with left (23°), right (23° or 11°), or dark lenses. We ended daily sessions with final trials using clear lenses. Upon successful landing on the target during a trial, we turned off the room lights to minimize short-term adaptation to prism lenses during prism trials and allowed bats a 2-minute rest period. Bats were offered a food reward during this rest period before the next trial.

In a subset of sessions, we attenuated auditory input during the flight task using bilateral earplugs (see *Earplugs and Hearing Tests* section) in combination with the prism lenses to manipulate both visual and auditory information. During earplug trials, the landing target was covered with green sound-absorbing material to further reduce echoes. We tested the ability of the bats to locate and land on the modified landing target during the preceding baseline trials (clear lenses) before beginning test trials (prism lenses + earplugs).

Each bat experienced all visual conditions in a counterbalanced order across days (within-subject, repeated measures). Sessions that included prism lenses and earplugs were counterbalanced to the nearest non-earplug visual condition for each bat. We tested each condition on ≥2 separate non-consecutive days to assess repeatability across days.

Audio-Video Recording Equipment

We recorded echolocation calls using a 32-channel array of electret condenser microphones (Pettersson Elektronik, Sweden). Each microphone's sensitivity and polar response were calibrated against a precision reference microphone (40DP 1/8", G.R.A.S., Denmark). Acoustic signals were band-pass filtered (10–100 kHz) at 10x gain (SBPBP-S1, Alligator Technology, CA) and digitized at 250 kHz. Three reflective markers rigidly mounted on the microphones defined the array coordinate frame for beamforming. We used a 13-camera Vicon system (T40/T40S; Oxford Metrics) to record three-dimensional flight trajectories at 200 fps (5 ms resolution). Four reflective markers on the helmet specified head pose during behavioral trials, permitting calculation of head-aim vectors. High-speed infrared video (Phantom Miro 310, 200 fps) provided corroborative behavioral footage for landing outcome scoring (see *Data Analysis* section).

Earplugs and Hearing Tests

In addition to the visual shift introduced by the prism lenses, we modified auditory input during behavioral trials using bilateral impression earmold plugs (Gold Velvet, Oklahoma City). Earmold plugs were inserted into each ear canal and molded to ensure a tight seal to attenuate incoming sound. Pinnae were glued down over the earplugs using spirit gum and the helmet was placed over the ears to further restrict pinna motion. We inspected the earplugs before and after each trial to ensure adequate placement in the ear canals for the duration of testing.

We assessed the effects of bilateral ear canal occlusion on hearing sensitivity by recording auditory brainstem responses (ABRs) from four bats (2 female, 2 male) with and without earplugs. Bats were anesthetized with 30 mg/kg ketamine and 4 mg/kg xylazine, administered intraperitoneally, and were placed in an IAC anechoic chamber on a warming pad to maintain body temperature at 37C. We recorded ABRs using subdermal electrodes (Rochester Horizon, Lifesync Neuro) placed at the ear, the vertex of the skull, and in the shoulder. Broadband clicks and tone pips from 4-58 kHz (5 ms, $\cos^2$ gated with 0.5 ms rise/fall) were broadcast by a speaker (MF1, Tucker Davis Technologies, Alachua, FL) placed 10 cm from the recording ear. Evoked potentials were recorded using a Medusa4Z preamplifier (TDT) connected to a TDT RZ6 processor and were filtered from 0.3-3 kHz using BioSigRZ software. For each bat, we measured ABRs with and without earplugs in a single recording session to ensure equivalent electrode placement across conditions.

Auditory detection thresholds were determined as the lowest sound level that generated an evoked response greater than 2 standard deviations above the noise floor and were visually verified. We compared within-subject threshold shifts between control and plug trials for clicks and tones to evaluate the ability of the earmold plugs to modify incoming acoustic information during behavioral trials. ABR thresholds were compared across control and plug trials using linear mixed-effects modeling, incorporating subject ID as a random effect for within-subject comparisons.

<u>Data Analysis</u>

Egyptian fruit bats emit brief, broadband echolocation clicks during flight, often producing clicks in closely spaced pairs. These click-pairs are frequently directed alternately to the left and right via rapid tongue movements, allowing the bat to scan space acoustically without large accompanying head rotations (*21*, *27*). As a result, the sonar emission direction can vary independently of head orientation, necessitating its direct estimation from acoustic data. We define the sonar beam pattern as a three-dimensional estimate of the spatial distribution of acoustic energy emitted by the bat for each click. Sonar beam patterns were reconstructed in MATLAB (*27*, *30*) by interpolating energy spectral density (ESD) across microphones. We computed maxima in the ESD maps to estimate beam aim for each click. Spherical spreading and atmospheric absorption were accounted for using daily temperature and humidity readings. We manually excluded clicks with poor quality ESD maps. Clicks where the beam aim pointed greater than 90° from the bat's heading were considered anomalous and excluded. Head direction was based on the vector passing through the front marker and the line connecting the back markers on the bat's helmet. When a marker was occluded, we used the bat's traveling direction to infer head direction.

All behavioral analysis was performed in 2D by projecting the bat's location, heading and beam aims onto the floor plane. Flights were segmented into trips by detecting changes in the bat's direction of motion away from the target: each trip captured one contiguous approach segment. Subsequent analysis was performed on the first outbound trip. Flight trajectories, head direction, and beam aim were synchronized in time. To calculate center of mass, each trajectory was first interpolated and resampled at equally spaced points to eliminate temporal information. Cumulative distance traveled was calculated by integrating the instantaneous speed of the bat over time. We estimated lateral deviation from the line connecting the release perch and the target landing perch and, in room coordinates, head direction (HD) as well as the bat's direction relative to the landing perch (target direction, TD). Head direction relative to target direction (HD-TD) was computed and median filtered with a 50 ms window. Click pairs were defined as pairs of consecutive clicks separated by < 40 ms. Unpaired clicks were excluded from the analysis. For each pair we computed the average beam direction to give the sonar direction (SD). Head direction relative to sonar direction (HD–SD) was computed by subtracting SD from the bat's HD in the interval between the two clicks of each pair, before boxcar filtering with a 300 ms window. We classified click-pairs with HD–SD < −3° as L-L and those with HD–SD > 3° as R-R. Sonar direction relative to target direction (SD-TD) was computed by subtracting the unwrapped, filtered HD-SD values from the unwrapped, filtered HD-TD values. The bat was considered to have approached the landing perch in a trial if, within the first trip, it spent > 100 ms within a 0.5 m radius of the target. We additionally computed IPI or inter-pair interval (boxcar filtered with a 300 ms window), flight speed (m/s), and tortuosity of the trajectory (cumulative distance traveled/displacement of the bat). We also manually classified trials as direct landings (DL), back-and-forth landings (BFL), back-and-forth failures (BFF) and direct failures (DF). A miss or collision with the landing perch was considered a failure. When averaging over a single trajectory, we unwrapped all circular data from -180° to 180° and computed circular means.

Binned analyses for HD-TD and lateral deviation of flight trajectories were performed by partitioning each trip into four cumulative-distance bins. Similarly, analyses for HD-SD and SD-TD, IPI, and flight speed were performed by partitioning each trip into six distance-to-target bins. We chose to do this for sonar-related measures because call timing is expected to depend on how close the bat is to the target rather than how far it has flown within a trip. For example, if a bat flew away from the target, we would not expect its click rate to continue increasing solely as a function of cumulative distance. We computed HD-SD or SD-TD far from the target and close to the target as the median value of click pairs between 1.5-2.5 m from the target and less than 1.5 m from the target respectively. When specified, behavioral variables were control-normalized by subtracting the average clear-lens measurement for that session.

To analyze learning related measures, we classified trials into "before" (two trials before prism exposure), "early" (first two trials after prism exposure), "late" (last two prism trials in each session) and "after" (two trials after prism exposure). Magnitude of learning per trial was determined as the average difference in lateral deviation between the late and early trials. Similarly, the magnitude of the after-effect was determined as the average difference between the after and before trials. To account for learning rates, we calculated the Z-scored lateral deviation by normalizing to the before-prism clear-lens trials ("baseline"). Trajectories were classified into left-shifted (Z-score > 1.5), right-shifted (Z-score < -1.5), and unshifted (-1.5 < Z-score < 1.5). We computed the number of trials after prism exposure until the bat executed a "corrected" trajectory as a proxy for the rate of learning, which was defined as non-left shifted for a left-shifting prism trial and non-right shifted for a right-shifting prism trial. All analysis was performed using custom scripts in Python.

Statistical analysis

We fit mixed-effects models to the behavioral data using custom scripts in Python. For continuous outcomes, we fit linear mixed-effects models with fixed effects of experimental condition, binned cumulative distance (for trajectory measures), binned distance to target (for sonar-related measures), trial block (pre- vs post-prism for after-effect analyses), and within-block trial number (trial number within a session for a given condition), together with the planned interactions experimental condition × distance (either cumulative or distance to target) and condition × within-block trial number. Random intercepts for bat, session, and trial number accounted for repeated observations across animals, day-to-day variation within individuals, and non-independence of multiple distance bins sampled from the same flight. Omnibus effects were evaluated with Wald $\chi^2$ tests, followed by post hoc contrasts from model-estimated marginal means when appropriate. Inter-pair interval was log-transformed to reduce skew; all other continuous variables were modeled on their native scales. Landing outcome was analyzed as a binary variable using a logistic generalized estimating equation with binomial link and exchangeable working correlation, clustered by session to account for repeated observations within a day. The model included condition, within-block trial number, and their interaction, with bat included as a covariate. Omnibus effects were evaluated with Wald $\chi^2$ tests, and planned contrasts comparing each condition with clear-lens control, as well as condition-specific slope differences relative to control, were Holm-corrected within the contrast family. To assess how bats performed under each condition, we also analyzed landing-type composition (DL, DF, BFF, BFL) using a global Pearson $\chi^2$ test, followed by planned pairwise $\chi^2$ comparisons against clear-lens controls with Holm correction; effect size was reported as Cramer's V.

Bayesian model

We modeled echolocating bats navigating under prism-induced auditory-visual conflict as ideal Bayesian observers maintaining a two-dimensional posterior probability density function $B(x, y, t)$ over target location (*50*). The posterior was updated multiplicatively every time the bat emits a click-pair by two likelihood channels $L_V(x, y, t)$ — a Gaussian centered at the prism-rotated apparent target — and an inter-aural level difference (ILD) likelihood $L_{ILD}(x, y, t, a)$ which depends on the sonar aim $a$: sharp on-axis and attenuated off-axis (Fig. S7, S8). The posterior was mixed with a uniform distribution based on its surprisal to mimic a "learning rate" (*50*). For action selection, the posterior was approximated by its modes: detecting peaks in $B$ yields discrete hypotheses $j$ with locations $x_j$ and probabilities $p_j$ (*50*). The bat's location was updated by a motor policy (*50*) that depended only on the current maximum-a-posteriori (MAP) estimate. Sonar aim was chosen in one of two ways before emitting each click-pair. In the "steering-coupled" policy, it pointed at the MAP estimate. In the "steering-decoupled" policy, it was selected by minimizing Bayes risk under a squared-error utility on the MAP estimate $R(a) = \sum_j p_j ||\hat{x}(a, x_j) - x_j||^2$, where $\hat{x}(a, x_j)$ was the posterior MAP that would result from taking action $a$ if hypothesis $j$ were true (*51*). Off axis attenuation of $L_{ILD}$ caused risk minimization to favor a policy that aimed the sonar towards secondary peaks in the posterior distribution (Supplementary Text — Theory).

### Supplementary Text — Theory

#### Overview

We sought to construct a Bayesian agent that could mimic the flight and sonar behaviors of the Egyptian fruit bat (*Rousettus aegyptiacus*) under auditory-visual conflict with the aim of exploring their adaptive significance. The state of the agent was specified by its 2D position and heading. The agent maintained a posterior belief about target location within a bounded environment, which was recursively updated by integrating auditory and visual evidence through their respective likelihood functions. The auditory likelihood function depended on the direction in which the bat aimed its sonar through a realistic model of *Rousettus* echolocation.

#### 1 Model of *Rousettus* Echolocation

The bat, located at the origin, emits a click-pair along its sonar aim, which is taken to be the reference direction for the echolocation model. Consider a stationary, focusing target located at $(R_T \cos\theta_T,\ R_T \sin\theta_T)$ with its focal point at the bat.[1] Its nonnegative real-valued reflectivity $\rho_{\theta_T}(\theta)$ depends on azimuth relative to the sonar aim $\theta$.

[1] This geometry eliminates inter-aural time and phase differences by construction.

At the target, the bat's click is measured as a complex-valued sound-pressure profile $s(t)$ scaled by a nonnegative real-valued beam $B(\theta)$. The pressure measured in the far field is given by the circular wave $p(t,\theta) = s(t)\,B(\theta)$. The received echo at the origin is then

$$e(t,\theta) = s\left(t - \frac{R_T}{c}\right) B(\theta)\,\rho_{\theta_T}(\theta). \tag{1}$$

where $c$ is the speed of sound. All acoustic transmission is assumed to be lossless. The head-related impulse response (HRIR) of the left ear is $h_l(t,\theta)$. Sound pressure at the left eardrum

$$p_l(t) = \iint e(\tau,\theta)\,h_l(t-\tau,\theta)\,d\tau\,d\theta \tag{2}$$

can be expressed in the frequency domain as

$$P_l(\omega) = \int E(\omega,\theta)\,H_l(\omega,\theta)\,d\theta, \tag{3}$$

where $E(\omega,\theta) = S(\omega)\,e^{-i\omega R_T/c}\,B(\theta)\,\rho_{\theta_T}(\theta)$ [2]. We assume a separable "head-related transfer function" (HRTF) $H_l(\omega,\theta) = G_{b,n_h}(\omega-\omega_0)\,\eta_l(\theta)\,e^{-i\omega d/c}$ for the left ear, where $d$ is the radius of the bat's head and $G_{b,n}$ is the Fourier transform of the Gammatone impulse response $t^{n-1}e^{-bt}\,\mathrm{H}(t)$ with bandwidth $b$ and shape parameter $n$. $\mathrm{H}(t)$ represents the Heaviside step function. We also assume that the HRTFs are coupled to the sonar aim of the bat, which could happen potentially through sonar-beam coupled movements of the ear-pinnae.

#### 1.1 Cochlear amplitude

By specifying the sound pressure profile and HRTFs, in addition to defining a cochlear filter, we can derive the amplitude experienced by the bat in each ear. For *Rousettus*, the click resembles a Gammatone $S(\omega) = G_{b,n_s}(\omega-\omega_0)$ matched to the bat's HRTF (Holland et al., 2004); $\omega_0/2\pi$ is the click's center frequency (CF), $b$ is the bandwidth and $n_s \approx n_h$ is the Gammatone shape parameter. The Gammatone product property[3] then gives

$$P_l(\omega) = \mathrm{B}(n_s,n_s)\,G_{b,2n_s}(\omega-\omega_0)\,e^{-i\omega(R_T+d)/c}\int \eta_l(\theta)\,B(\theta)\,\rho_{\theta_T}(\theta)\,d\theta. \tag{4}$$

A cochlear filter matched to the click's CF filters the sound pressure at the ear. In the frequency domain, this multiplies (4) by $K_{\omega_0}(\omega) = G_{b,n_s}(\omega-\omega_0)$, giving

$$\Psi_l(\omega) = \mathrm{B}(n_s,2n_s)\,\mathrm{B}(n_s,n_s)\,G_{b,3n_s}(\omega-\omega_0)\,e^{-i\omega(R_T+d)/c}\int \eta_l(\theta)\,B(\theta)\,\rho_{\theta_T}(\theta)\,d\theta. \tag{5}$$

Since $G_{b,3n_s}$ is the Fourier transform of the Gammatone $t^{3n_s-1}\,e^{-bt}\,e^{-i\omega_0 t}\,\mathrm{H}(t)$, the instantaneous envelope of the cochlear-filtered waveform is

$$A_l(t) = \mathrm{B}(n_s,2n_s)\,\mathrm{B}(n_s,n_s)\left[t - \tfrac{R_T+d}{c}\right]^{3n_s-1} e^{-b\,[t-(R_T+d)/c]}\,\mathrm{H}\!\left(t - \tfrac{R_T+d}{c}\right)\int \eta_l(\theta)\,B(\theta)\,\rho_{\theta_T}(\theta)\,d\theta. \tag{6}$$

#### 1.2 Inter-aural level difference (ILD)

Comparing amplitude of the echo between the right and left ear allows the bat to estimate the angular position of the target. The ILD for click $k$ in dB is the average log-amplitude ratio over a window $w(t) = \mathrm{H}(t) - \mathrm{H}(t-(R_M+d)/c)$ that opens at click emission and closes after $(R_M+d)/c$ seconds. $R_M$ is the maximum target range the bat waits to hear back from:

$$\mathrm{ILD}_k \equiv \frac{20}{\log 10}\,\frac{\int w(t)\,\log\!\left(A_r^{(k)}(t)/A_l^{(k)}(t)\right)dt}{\int w(t)\,dt} = \frac{20}{\log 10}\log\frac{\int \eta_r(\theta)\,B_k(\theta)\,\rho_{\theta_T}(\theta)\,d\theta}{\int \eta_l(\theta)\,B_k(\theta)\,\rho_{\theta_T}(\theta)\,d\theta} \tag{7}$$

[2] $S(\omega)$ is the Fourier transformed sound pressure profile $s(t)$

[3] $G_{b,m}(\omega-\omega_0)\,G_{b,n}(\omega-\omega_0) = \mathrm{B}(m,n)\,G_{b,m+n}(\omega-\omega_0)$, where $\mathrm{B}(m,n) = \Gamma(m)\Gamma(n)/\Gamma(m+n)$ is the Beta function.

#### 1.3 Beam patterns

The bat emits a "right-left" click-pair. Each click $k \in \{0, 1\}$ is modeled as a shifted circular normal[4] beam with width $\sigma_B$, half the inter-pair angle $\tilde{\phi}$ and maximum amplitude $C$,

$$B_k(\theta) = C\, \mathcal{V}_{\sigma_B}\big(\theta;\, (-1)^k\, \tilde{\phi}\big), \qquad \sigma_B = \max(\sigma_*, \sigma_T) \tag{8}$$

with $\kappa_B = 1/\sigma_B^2$ and $\kappa_T = 1/\sigma_T^2$. $\tilde{\phi}$ depends on the target angular size $\sigma_T$ through a single parameter $\sigma_*$.

$$\tilde{\phi} = \frac{x_0^3\, \sigma_*^3}{x_0^2\, \sigma_*^2 + \sigma_T^2}, \tag{9}$$

We chose this form since the inter-pair angle decreases as the bat gets close to the target (Yovel et al., 2010). To constrain $x_0$, we note that when the bat is far from the target and aims at it, the maximum slope of the beam lands on the target (Yovel et al., 2010); i.e., when the target is small, we need the beam width to equal half the inter-pair angle $\sigma_B = \tilde{\phi}$. When $\sigma_T \ll \sigma_*$, we have $\sigma_B = \sigma_*$ (8) and $\tilde{\phi} \to x_0\, \sigma_*$ (9) implying $x_0$ is the solution to $x^3 - x^2 - 1 = 0$. When the target is large ($\sigma_T \gg x_0\, \sigma_*$), $\sigma_B = \sigma_T$, $\tilde{\phi} \to 0$ and the beams overlap. $C$ is chosen so the on-axis integrated loudness $\int B_k\, T\, d\theta'$ is distance independant, which is consistent with bats producing softer clicks the closer they are to the target.

$$C \;=\; \frac{I_0(\kappa_B)\, I_0(\kappa_T)}{I_0(\kappa_B + \kappa_T)}, \tag{10}$$

Note that we have assumed the bat needs to know the instantaneous angular size of the target before it emits a click. This can likely be estimated from the visual information available to the bat.

#### 1.4 1D ILD likelihood

Given an instantaneous $\mathrm{ILD}(\theta_T)$ measurement, the bat can calculate the likelihood of the target being at angle $\theta$. We assumed ILD measurements are subjected to logistic noise with scale $s_{\mathrm{ILD}}$ in dB. For a single click $k$, the likelihood function is then

$$\mathcal{L}_k(\theta) = \mathrm{sech}^2\left(\frac{\mathrm{ILD}_k(\theta_T) - \mathrm{ILD}_k(\theta)}{2 s_{\mathrm{ILD}}}\right). \tag{11}$$

We note that a biologically realistic circuit model can compute a likelihood of this form via a spatial derivative of a sigmoidal population code (Spence & Pearson, 1989). Setting $s_{\mathrm{ILD}} = 20/\log 10 \approx 8.7\,\mathrm{dB}$, we define

$$f_k(\theta) \equiv \frac{\int [\eta_r(\theta') - \eta_l(\theta')]\, B_k(\theta')\, \rho_\theta(\theta')\, d\theta'}{\int [\eta_r(\theta') + \eta_l(\theta')]\, B_k(\theta')\, \rho_\theta(\theta')\, d\theta'} \tag{12}$$

such that $\tanh(\mathrm{ILD}_k(\theta)/2s_{\mathrm{ILD}}) = f_k(\theta)$. The numerator resembles a linear pressure-difference filter (Altes, 1978). Writing (11) in terms of $f_k$,

$$\mathcal{L}_k(\theta) = \frac{\big(1 - f_k(\theta_T)^2\big)\big(1 - f_k(\theta)^2\big)}{\big(1 - f_k(\theta_T)\, f_k(\theta)\big)^2}. \tag{13}$$

Following HRTF measurements in *Rousettus* (Nikolić et al., 2012), we pick $\eta_{r,l}(\theta) = \cos^2\big((\theta \mp \theta_H)/2\big)$ with $\theta_H = \pi/2$, implying the bat is most sensitive to sounds exactly to its right and left. This gives, $\eta_r + \eta_l = 1$ and $\eta_r - \eta_l = \sin\theta$, so the numerator and denominator of (12) become $\int \sin(\theta')\, B_k\, \rho_\theta\, d\theta'$ and $\int B_k\, \rho_\theta\, d\theta'$ respectively.

For click $k = 0$, since $B_0(\theta') = C\mathcal{V}_{\sigma_B}(\theta'; \tilde{\phi})$ and $\rho_\theta(\theta') = \mathcal{V}_{\sigma_T}(\theta'; \theta)$, using the product identity [5] we have

$$C\mathcal{V}_{\sigma_B}(\theta'; \tilde{\phi})\, \mathcal{V}_{\sigma_T}(\theta'; \theta) = \frac{C I_0(1/\sigma^2)}{I_0(1/\sigma_B^2)\, I_0(1/\sigma_T^2)}\, \mathcal{V}_\sigma(\theta'; \mu), \tag{14}$$

[4] The *circular normal* (von Mises) distribution $\mathcal{V}_\sigma(\theta; \mu) = \exp\big(\cos(\theta - \mu)/\sigma^2\big) / \big(2\pi I_0(1/\sigma^2)\big)$ is a Gaussian-like density on the circle with mean direction $\mu$ and angular width $\sigma$ (concentration $1/\sigma^2$); $I_n$ is the modified Bessel function of the first kind of order $n$.

[5] $\mathcal{V}_{\sigma_1}(\theta; \mu_1)\, \mathcal{V}_{\sigma_2}(\theta; \mu_2) = \dfrac{I_0(1/\sigma^2)}{I_0(1/\sigma_1^2)\, I_0(1/\sigma_2^2)}\, \mathcal{V}_\sigma(\theta; \mu)$, where $(\sigma, \mu)$ satisfy $\dfrac{1}{\sigma^2} e^{i\mu} = \dfrac{1}{\sigma_1^2} e^{i\mu_1} + \dfrac{1}{\sigma_2^2} e^{i\mu_2}$.

where $\frac{1}{\sigma^2} e^{i\mu} = \frac{1}{\sigma_B^2} e^{i\tilde{\phi}} + \frac{1}{\sigma_T^2} e^{i\theta}$. Substituting (14) into the denominator of (12):

$$\int C\mathcal{V}_{\sigma_B}(\theta'; \tilde{\phi})\, \mathcal{V}_{\sigma_T}(\theta'; \theta)\, d\theta' = \frac{C I_0(1/\sigma^2)}{I_0(1/\sigma_B^2)\, I_0(1/\sigma_T^2)} \tag{15}$$

since the circular normal integrates to 1. Substituting (14) into the numerator of (12):

$$\int \sin(\theta')\, C\mathcal{V}_{\sigma_B}(\theta'; \tilde{\phi})\, \mathcal{V}_{\sigma_T}(\theta'; \theta)\, d\theta' = \frac{C I_0(1/\sigma^2)}{I_0(1/\sigma_B^2)\, I_0(1/\sigma_T^2)} \frac{I_1(1/\sigma^2)}{I_0(1/\sigma^2)} \sin(\mu) \tag{16}$$

from the first sine moment of the circular normal. Dividing (16) by (15),

$$f_0(\theta) = \frac{I_1(1/\sigma^2)}{I_0(1/\sigma^2)} \sin(\mu), \tag{17}$$

By the reflection symmetry of the beam pair about the midline,

$$f_1(\theta) = -f_0(-\theta). \tag{18}$$

Since each click yields an independent likelihood (13), the fused 1D likelihood from a click-pair is their product. From (18):

$$\mathcal{L}_{\text{1D}}(\theta) = \prod_{\epsilon=\pm1} \frac{\left(1 - f_0(\epsilon\, \theta_T)^2\right)\left(1 - f_0(\epsilon\, \theta)^2\right)}{\left(1 - f_0(\epsilon\, \theta_T)\, f_0(\epsilon\, \theta)\right)^2}. \tag{19}$$

### 2 Bayesian Agent

We adapt a model of rodent navigation from AbdelRahman et al. (2026) for the bat in our task. It consists of a Bayesian agent with the following components.

#### 2.1 State, Belief and Policies

The agent's state at time $t$ consists of 2D position $\mathbf{b}_t \in \mathbb{R}^2$ and heading $\psi_t \in [-\pi, \pi)$. The posterior belief $B(\mathbf{x}, t)$ over the target location is discretized on a uniform grid $\Omega \subset \mathbb{R}^2$ of size $N \times N$. Each grid point $\mathbf{x} = (x_x, x_y) \in \Omega$ subtends an angle $\theta(\mathbf{x}; \mathbf{b}_t)$ relative to the bat. The agent attempts to intercept a target placed at $\mathbf{x}_T \in \mathbb{R}^2$ by controlling $\mathbf{b}_t$ and $\psi_t$ via a flight policy and sonar aim $\beta \in [-\pi, \pi)$, which specifies the direction of the click-pair, via a sonar policy.

#### 2.2 2D likelihoods

Each sensory channel contributes a two-dimensional likelihood function $\mathcal{L}(\mathbf{x}, t)$ over the grid $\Omega$. The likelihood scores the plausibility of location $\mathbf{x}$ containing the true target, given the current sensory measurement. Similar to AbdelRahman et al. (2026) we scale the likelihood to lie between $l$ and $1 - l$

$$\mathcal{L}(\mathbf{x}, t) = \ell + (1 - 2\ell)\, \frac{\widetilde{\mathcal{L}}(\mathbf{x}, t)}{\max\limits_{\mathbf{x}' \in \mathbb{R}^2} \widetilde{\mathcal{L}}(\mathbf{x}', t)}. \tag{20}$$

to prevent the belief from being zeroed out anywhere on the grid, which precludes all future updates. We normalize by the global maximum over $\mathbb{R}^2$ rather than over the grid $\Omega$ to avoid edge effects introduced by the bounded environment.

##### 2.2.1 Visual likelihood

The passive visual channel is a circular Gaussian centered at the apparent target position $\mathbf{x}_{\text{app}}$,

$$\widetilde{\mathcal{L}_V}(\mathbf{x}, t) \;=\; \exp\!\left(-\frac{\left\|\mathbf{x} - \mathbf{x}_{\text{app}}\right\|^2}{2\,\sigma_{\text{vis}}^2}\right), \tag{21}$$

normalized (20) to give $\mathcal{L}_V(\mathbf{x}, t)$. A prism manipulation rotates $\mathbf{x}_{\text{app}}$ relative to the true target by an angle $\Delta$ about the bat: $\mathbf{x}_{\text{app}} = R(\Delta)(\mathbf{x}_T - \mathbf{b}_t) + \mathbf{b}_t$, where $R(\Delta)$ is a $2 \times 2$ rotation matrix. Note that the apparent allocentric position of the visual target depends on the distance of the bat from the true target.

###### 2.2.2 ILD likelihood

We construct $\mathcal{L}_{\text{ILD}}$ from the $\mathcal{L}_{1\text{D}}$ (19) in four steps.

**(i) Compute bat–target geometry using $(\mathbf{x}_T, \mathbf{b}_t, \beta)$.** We have

$$\theta_T = \operatorname{atan2}\left(x_T - b_{t,x},\, y_T - b_{t,y}\right) - \beta, \qquad \sigma_T = r_T / \|\mathbf{x}_T - \mathbf{b}_t\|, \qquad r = \|\mathbf{x}_T - \mathbf{b}_t\| \tag{22}$$

where $r_T$ is the target radius (m). $\sigma_T$ is estimated instantaneously by the visual system. The click-pair parameters $(C, \sigma_B, \tilde{\phi})$ then follow from (8)–(10).

**(ii) Find peaks in the 1D click-pair likelihood.** Numerically compute

$$\{\hat{\theta}_k\} \;=\; \arg\max_{\theta}\, \mathcal{L}_{1\text{D}}(\theta \mid \theta_T) \tag{23}$$

by sampling $\mathcal{L}_{1\text{D}}$ from (19) on a ring of $N_{\text{ring}}$ angles around the bat.

**(iii) Compute the echo amplitude.** The total sound pressure due to the click-pair is the sum of the integrated loudness $D_k(\theta_T) \equiv \int B_k(\theta')\, \rho_{\theta_T}(\theta')\, d\theta'$ for clicks $k \in \{0, 1\}$. Using the product identity (14),

$$A_{\text{echo}}(\theta_T; \sigma_B, \sigma_T) \;=\; D_0(\theta_T) + D_1(\theta_T), \quad D_k(\theta_T) \;=\; \frac{I_0\Big(\kappa_p(\theta_T - (-1)^k\, \tilde{\phi})\Big)}{I_0(\kappa_B + \kappa_T)}, \tag{24}$$

where $\kappa_T = 1/\sigma_T^2$ and $\kappa_p(\theta) = \sqrt{\kappa_B^2 + \kappa_T^2 + 2\,\kappa_B\,\kappa_T\,\cos\theta}$.

**(iv) Place amplitude-dependent circular Gaussians at peaks.** Each peak $\hat{\theta}_k$ contributes a circular Gaussian centered at distance $r$ from the bat, with spatial width $\sigma_{2D} = \sigma_{\text{ILD}} / A_{\text{echo}}(\theta_T)$. Well-aimed click-pairs yield sharp localized Gaussians, while poorly-aimed pairs yield flat, weakly informative ones.

The raw 2D likelihood is a sum of unit-height Gaussians:

$$\widetilde{\mathcal{L}_{\text{ILD}}}(\mathbf{x}) \;=\; \sum_k \exp\left(-\frac{\|\mathbf{x} - \mathbf{c}_k\|^2}{2\,\sigma_{2D}^2}\right), \quad \mathbf{c}_k = \mathbf{b}_t + r \begin{pmatrix} \sin(\beta + \hat{\theta}_k) \\ \cos(\beta + \hat{\theta}_k) \end{pmatrix}, \tag{25}$$

All Gaussians share width $\sigma_{2D}$. Since the maximum value of $\widetilde{\mathcal{L}_{\text{ILD}}}$ is close to 1 over $\mathbb{R}^2$, the normalized $\mathcal{L}_{\text{ILD}}(\mathbf{x}) \approx 1\ \forall\ \mathbf{x} \in \Omega$. This means that a soft echo does not modify the belief strongly.

##### 2.3 Belief update with surprisal gating

Every $\tau_b$ seconds, the bat performs a multiplicative Bayes update. Then the posterior is mixed with a uniform distribution weighted by its surprisal (AbdelRahman et al., 2026):

$$\widetilde{B}_{t+\tau_b}(\mathbf{x}) \;\propto\; B_t(\mathbf{x})\, \mathcal{L}_V(\mathbf{x})\, \mathcal{L}_{\text{ILD}}(\mathbf{x} \mid \mathbf{x}_T, \mathbf{b}_t, \beta_t), \tag{26}$$

$$B_{t+\tau_b}(\mathbf{x}) \;=\; (1 - \alpha_t)\, \widetilde{B}_{t+\tau_b}(\mathbf{x}) + \alpha_t\, u(\mathbf{x}), \tag{27}$$

$$\alpha_t \;=\; \frac{1}{1 + e^{-(\mathcal{S}_t - \mathcal{S}_0)/\gamma}} \tag{28}$$

where $u(\mathbf{x}) = 1/|\Omega|$ is the uniform distribution, $\mathcal{S}_t = D_{\text{KL}}\big(\widetilde{B}_{t+\tau_b} \,\big\|\, B_t\big)$ is the Bayesian surprise (surprisal) and $(\mathcal{S}_0, \gamma)$ parametrize the gating threshold and slope. High surprisal flattens the belief whenever new evidence is sharply inconsistent with the current belief. Our model has a low threshold to reduce the influence of spatial memory on the estimate of target location.

#### 2.4 Discrete hypothesis space

Following AbdelRahman et al. (2026), we approximate the continuous posterior by its modes: peak-finding on $B_t$ yields a discrete hypothesis space $\{\mathbf{x}_j\}_{j=0}^{K-1}$ of candidate target locations with probabilities

$$p_j \;\propto\; B_t(\mathbf{x}_j), \qquad \sum_j p_j = 1, \tag{29}$$

i.e. $p_j$ is the (normalized) height of belief peak $j$.

#### 2.5 Flight policy

The bat commits to flying toward the maximum-a-posteriori (MAP) estimate of its belief —

$$\hat{\mathbf{x}}_t \;=\; \arg\max_{\mathbf{x}\in\Omega} B_t(\mathbf{x}). \tag{30}$$

The flight target $\hat{\mathbf{x}}_t$ is recomputed at a planning timescale $\tau_{\text{plan}}$ larger than the belief-update timescale $\tau_b$ such that the bat moves at a constant linear speed $v$ and, between planning events, advances along the unique circular arc that passes through $\mathbf{b}_t$ and $\hat{\mathbf{x}}_t$ by integrating

$$\dot{\mathbf{b}}_t = v\,\hat{e}(\psi_t), \qquad \dot{\psi}_t = \omega_t, \tag{31}$$

where $\psi_t$ is the heading and $\hat{e}(\psi) = (\cos\psi, \sin\psi)$ is the heading unit vector. The angular velocity $\omega_t$ is chosen at each planning event to arc the bat toward $\hat{\mathbf{x}}_t$:

$$\omega_t \;=\; \begin{cases} \dfrac{2\,v\,\sin\delta_t}{\|\hat{\mathbf{x}}_t - \mathbf{b}_t\|} & \text{if } |\delta_t| \le \pi/2 \quad \text{(when target is ahead)} \\ \dfrac{2\,v\,\delta_t}{\|\hat{\mathbf{x}}_t - \mathbf{b}_t\|} & \text{otherwise} \quad \text{(when target is behind)} \end{cases} \tag{32}$$

where $\delta_t = \operatorname{atan2}\left(\hat{x}_{t,y} - b_{t,y},\, \hat{x}_{t,x} - b_{t,x}\right) - \psi_t$ is the heading error (wrapped to $[-\pi, \pi)$). When the target is behind, switching to a proportional control policy prevents the bat from reducing its rotational velocity when $\delta_t$ is greatest but $\sin\delta_t$ is 0.

#### 2.6 Sonar policy

The sonar aim in the **steering-coupled policy** tracks the MAP estimate (30):

$$\beta_t \;=\; \theta(\hat{\mathbf{x}}_t;\, \mathbf{b}_t) \;=\; \operatorname{atan2}\big(\hat{x}_{t,x} - b_{t,x},\; \hat{x}_{t,y} - b_{t,y}\big). \tag{33}$$

The sonar aim in the **steering-decoupled policy** is chosen by minimizing the Bayes risk of a squared-error utility on the MAP estimate (Müller, 1999). The bat chooses a sonar aim $\beta \in \{\beta^{(0)}, \dots, \beta^{(K-1)}\}$ pointing at one of the $K$ belief peaks, $\beta^{(a)} = \theta(\mathbf{x}_j[a]; \mathbf{b}_t)$. For each candidate action $a$ and hypothesis $\mathbf{x}_j$, the simulated posterior (Müller, 1999) under a noiseless echo from $\mathbf{x}_j$ and beam $\beta^{(a)}$ is

$$\widetilde{B}'(\mathbf{x} \mid a, j) \;\propto\; B_t(\mathbf{x})\,\mathcal{L}_{\text{ILD}}\Big(\mathbf{x} \mid \mathbf{x}_j, \mathbf{b}_t, \beta^{(a)}\Big), \tag{34}$$

with MAP estimate

$$\hat{\mathbf{x}}(a, \mathbf{x}_j) \;=\; \arg\max_{\mathbf{x}\in\Omega} \widetilde{B}'(\mathbf{x} \mid a, j). \tag{35}$$

The utility is then

$$\mathcal{J}[j, a] \;=\; \|\hat{\mathbf{x}}(a, \mathbf{x}_j) - \mathbf{x}_j\|^2, \tag{36}$$

and the Bayes risk (expected utility under the bat's belief) is

$$R(a) \;=\; \sum_{j=0}^{K-1} p_j\, \mathcal{J}[j, a]. \tag{37}$$

The chosen action is

$$\boxed{a_t^* \;=\; \arg\min_{a\in\{0,\ldots,K-1\}} R(a).} \tag{38}$$

Consider $K = 2$ distant peaks at $\mathbf{x}_0, \mathbf{x}_1$ with probabilities $p_0, p_1$. From the squared-error utility, we get:

$$\begin{aligned} R(0) &= p_0\,\|\hat{\mathbf{x}}(0,\mathbf{x}_0) - \mathbf{x}_0\|^2 + p_1\,\|\hat{\mathbf{x}}(0,\mathbf{x}_1) - \mathbf{x}_1\|^2, \\ R(1) &= p_0\,\|\hat{\mathbf{x}}(1,\mathbf{x}_0) - \mathbf{x}_0\|^2 + p_1\,\|\hat{\mathbf{x}}(1,\mathbf{x}_1) - \mathbf{x}_1\|^2. \end{aligned} \tag{39}$$

Suppose $p_1 > p_0$, then the MAP estimate is at $\mathbf{x}_1$. Probing peak $j$ confirms it on-axis: $\hat{\mathbf{x}}(j, \mathbf{x}_j) = \mathbf{x}_j$. Probing peak $a \neq j$ leaves the MAP estimate at $\mathbf{x}_1$: $\hat{\mathbf{x}}(a, \mathbf{x}_j) = \mathbf{x}_1$. Substituting,

$$\begin{aligned} R(0) &= p_0\,\|\mathbf{x}_0 - \mathbf{x}_0\|^2 + p_1\,\|\mathbf{x}_1 - \mathbf{x}_1\|^2 = 0, \\ R(1) &= p_0\,\|\mathbf{x}_1 - \mathbf{x}_0\|^2 + p_1\,\|\mathbf{x}_1 - \mathbf{x}_1\|^2 = p_0\,d^2 \end{aligned}$$

where $d = \|\mathbf{x}_0 - \mathbf{x}_1\|$. Hence $R(0) < R(1)$ and $a^* = 0$: the bat probes the lower-probability peak.

The intuition for this decision as follows. The bat always has the option of probing either the visual or the auditory target. Suppose the bat wrongly believes the target location is specified by vision. Probing the visual target would retain its belief in vision given the manner in which acoustic evidence is integrated since pointing off-axis doesn't update the belief. On the other hand probing the auditory target would shift the belief to the correct location. Therefore, it's riskier to continue aiming at the visual target even if you currently believe in it.

#### 2.7 Parameter values

Table 1 lists the parameter values used in the simulations reported in the main text.

#### 2.8 Summary

The bat (i) flies towards the MAP estimate (31)–(32), (ii) selects $a^*$ via (38), (iii) updates $B$ via (26)–(28) and the cycle repeats.

Table 1: Simulation parameters.

| Symbol | Description | Value | Units |
|---|---|---|---|
| *Grid* | | | |
| $\Omega$ | spatial extent | $[-2.5, 2.5] \times [0, 5]$ | m |
| $N_x \times N_y$ | belief-grid size | $200 \times 200$ | cells |
| $N_{\text{ring}}$ | angular grid | 720 | cells |
| *Target* | | | |
| $\mathbf{x}_T$ | true target position | (0, 3.5) | m |
| $r_T$ | target radius | 0.25 | m |
| *Initial state* | | | |
| $\mathbf{b}_0$ | bat initial position | (0, 0) | m |
| $\psi_0$ | bat initial heading | 90 | deg |
| $\Delta$ | prism rotation | 23 | deg |
| $v$ | flight speed | 1 | m/s |
| *Initial belief (mixture of Gaussians)* | | | |
| $(w_V, w_A)$ | weights (visual, auditory) | (0.6, 0.4) | — |
| $\sigma_{\text{vis}}$ | visual prior and likelihood width | 0.2 | m |
| $\sigma_{\text{aud}}$ | auditory prior width | 0.2 | m |
| *Timescales* | | | |
| $\tau_b$ | belief-update interval | 0.08 | s |
| $\tau_{\text{plan}}$ | flight replan interval | 0.5 | s |
| $T_{\max}$ | total simulation time | 4.5 | s |
| *Surprisal gate* | | | |
| $\mathcal{S}_0$ | surprisal threshold | 5.5 | nats |
| $\gamma$ | sigmoid sharpness | 1.0 | $\text{nats}^{-1}$ |
| *Echolocation* | | | |
| $\sigma_*$ | beam max-slope parameter | 2.5 | deg |
| $\sigma_{\text{ILD}}$ | on-axis ILD Gaussian width | 0.2 | m |
| *Likelihood* | | | |
| $\ell$ | likelihood floor | 0.05 | — |

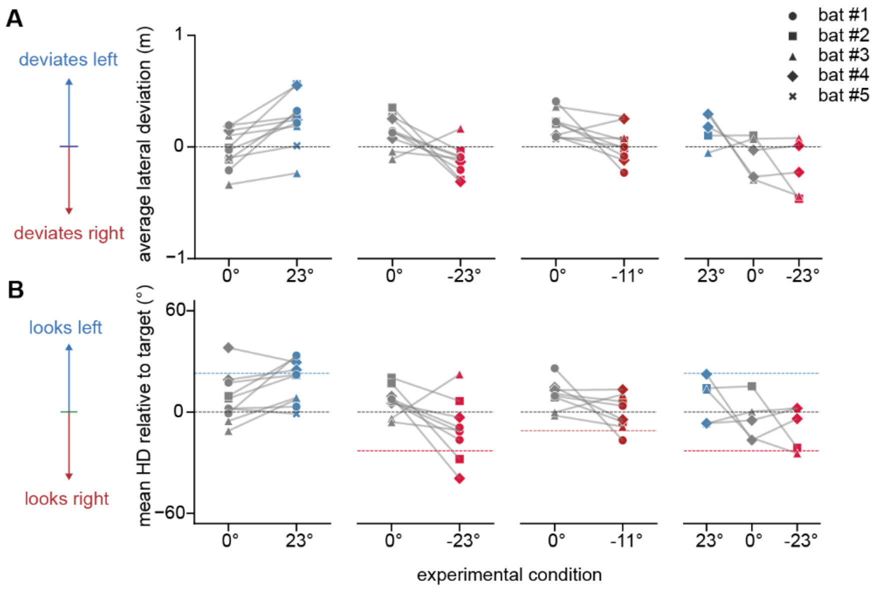

**Fig S1. Smaller prism-shifts produce a smaller but directionally consistent trajectory and head direction bias.**
A) Average lateral flight deviation and B) head direction relative to the stationary target shown for each bat across cumulative distance flown for the larger left and right prism conditions (23°), smaller right prism condition (11°), and combined left and right larger prism conditions.

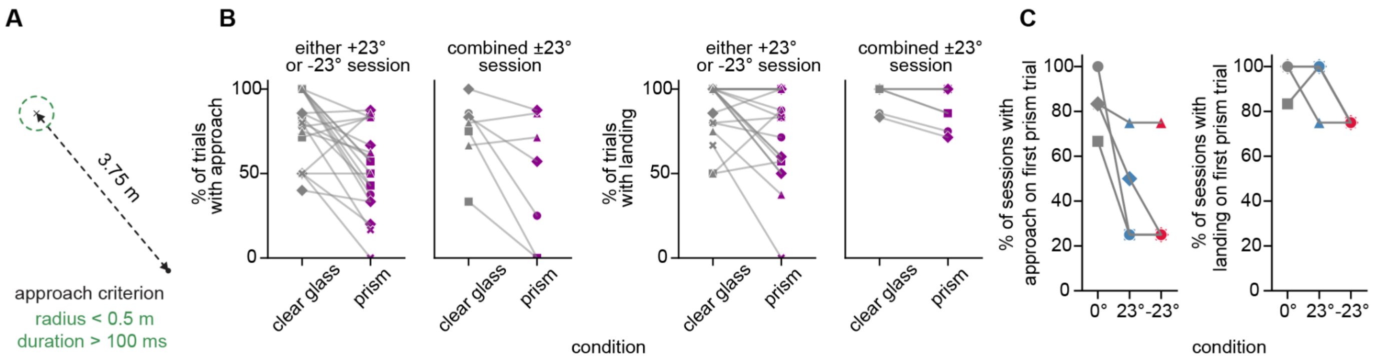

**Fig S2. Bats can course correct with prism lenses.**

A) Approach criterion: bat entered a 0.5 m radius around the target for ≥100 ms during the first trip of the trial. B) Direct landings decreased under large prism shifts compared to clear lenses. C) Percent of sessions in which bats approached the target on the first trial. Symbols represent data from different bats.

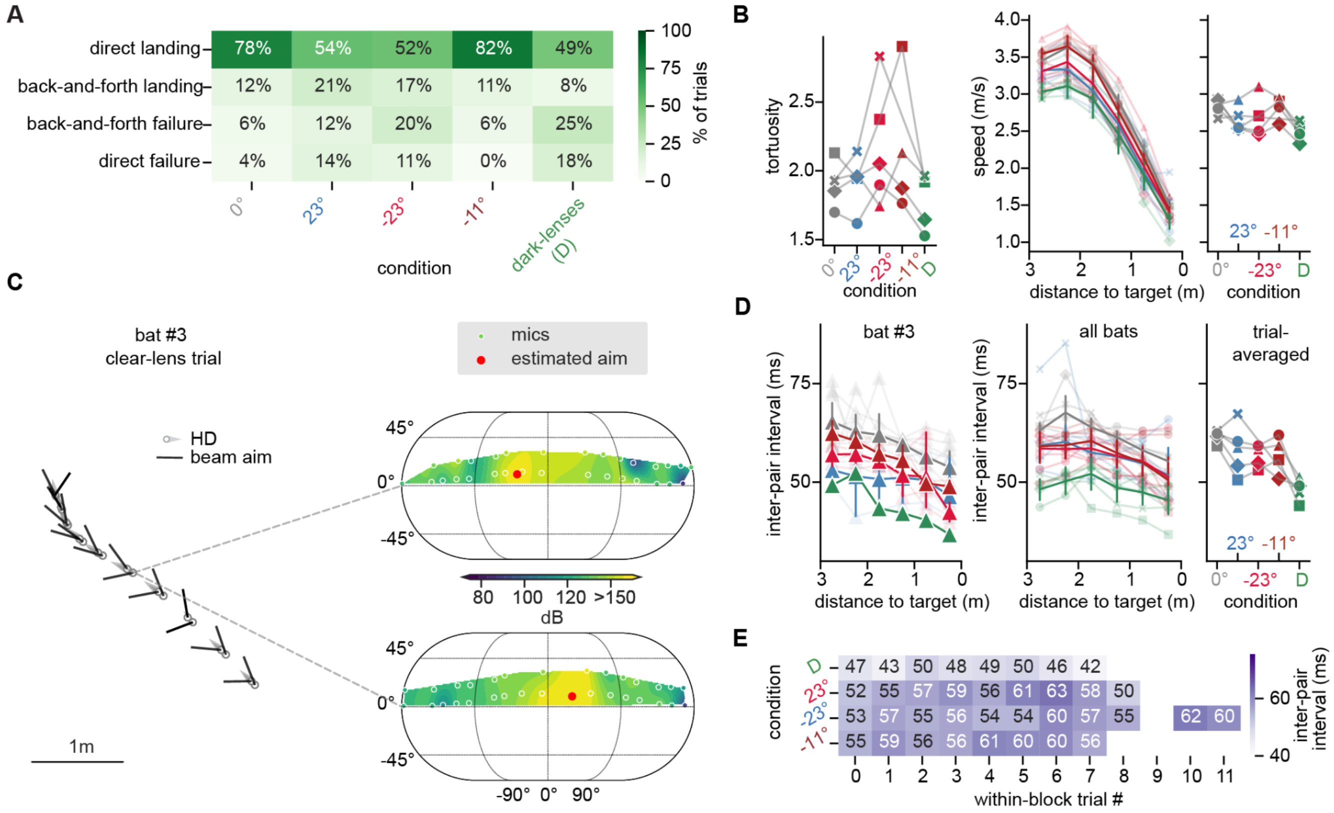

**Fig S3. Steering and biosonar depend on the availability and unreliability of vision.**

A) Landing outcomes across conditions with visual perturbations. B) (Left) Tortuosity of the flight path measured as distance traveled/displacement across conditions. (Middle) Flight speed as a function of distance to target averaged over trials and bats across conditions. (Right) Flight speed averaged across all trials for each bat. C) Example clear-lens flight trajectory showing the typical alternating left–right click pattern, with the power spectral density associated with each click. D) (Left) Inter-pair interval across distance to target for a representative session from bat #3, (Middle) pooled data across all bats, and (Right) trial-averaged values for clear lenses, large left-shifting prisms, large right-shifting prisms, smaller right-shifting prisms, and dark lenses. Shorter intervals indicate increased click rate during approach and under conditions in which visual information was unavailable or spatially displaced. E) Mean inter-pair interval for each within-block trial, obtained by averaging over trials and then across bats. Symbols represent data from different bats.

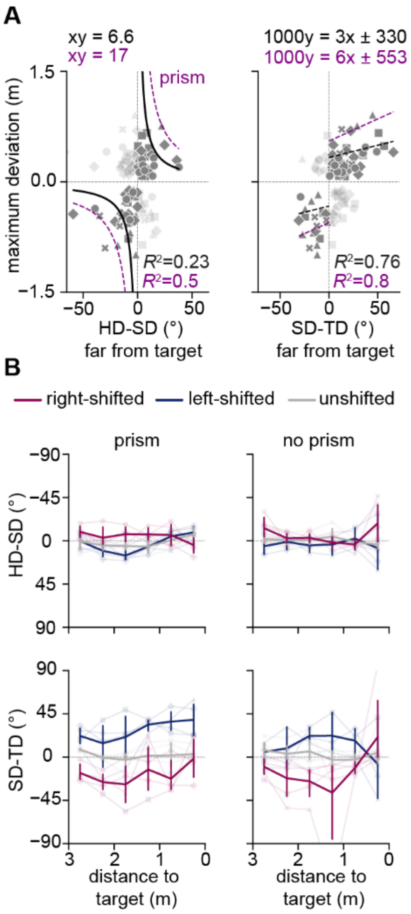

**Fig S4. Biosonar does not diverge in non-prism trials.**

A) Greater early head–sonar divergence (left) was less strongly associated with smaller maximum trajectory deviation with clear-lenses than prisms (lower $R^2$), suggesting that sampling away from the current heading was not required for approaching the target. Similarly, early aiming at the target was less predictive of maximum trajectory deviation for clear lenses than for prisms, as indicated by the approximately twofold lower slope. B) Control-normalized head-sonar divergence (top) and sonar direction relative to target (bottom) across distance to the target with and without prisms for right-, left- and unshifted trials showing that HD-SD divergence is prominent with but not without prisms for similar trajectories. Note that SD-TD divergence can be achieved even without HD-SD divergence purely due to deviations in the trajectory. Symbols represent data from different bats.

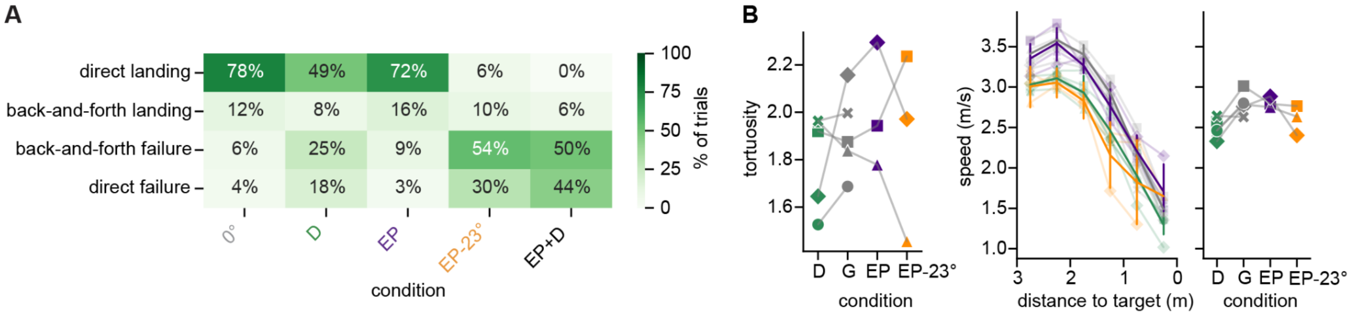

**Fig S5. Earplugs increase failure rate when vision is unavailable or unreliable.**

A) Landing outcomes across conditions with auditory perturbations. B) (Left) Tortuosity of the flight path measured as distance traveled/displacement across conditions. (Middle) Flight speed as a function of distance to target averaged over trials and bats across conditions. (Right) Flight speed averaged across all trials for each bat. Symbols represent data from different bats.

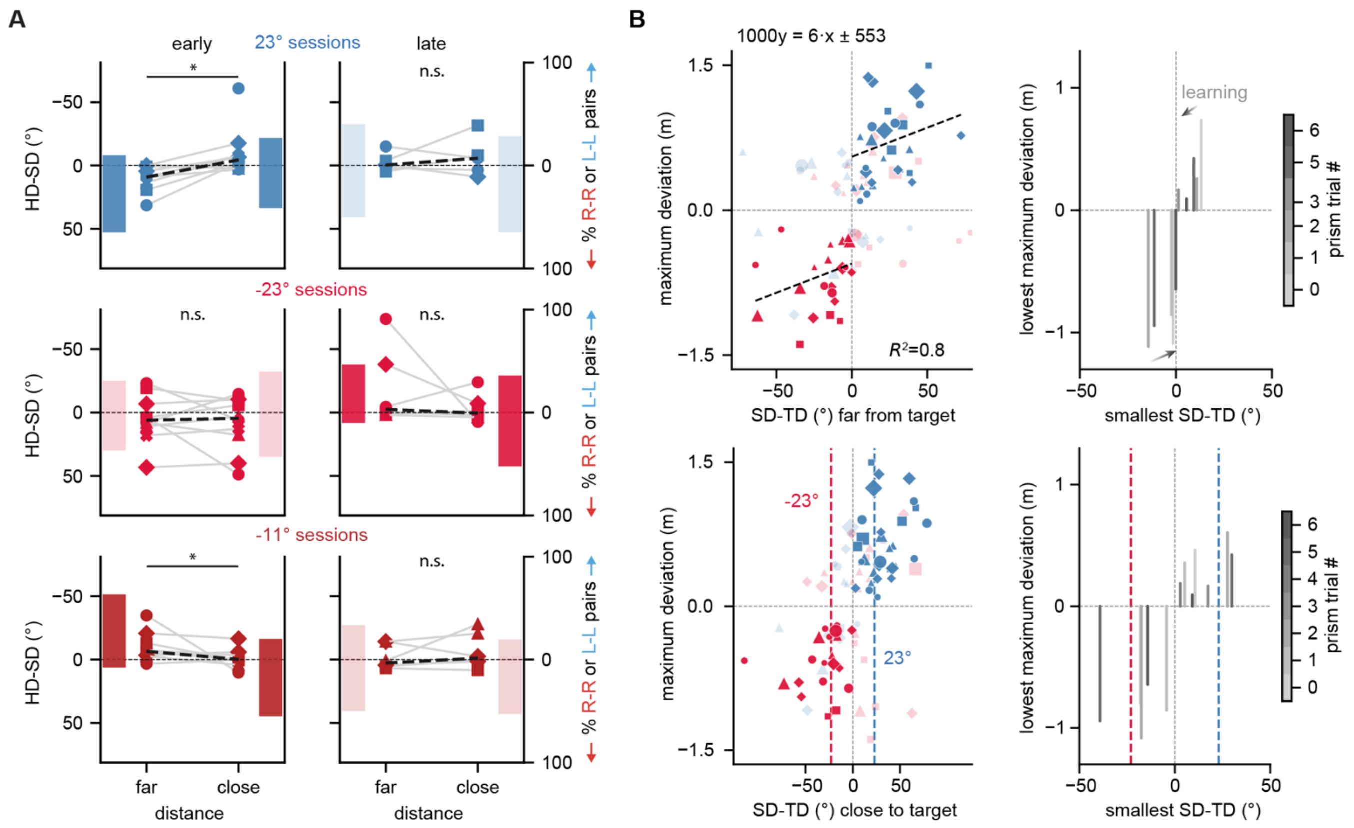

**Fig S6. Sonar becomes more consistent over trials.**

A) Head–sonar divergence far from and close to the target during early and late left (top), large right (middle), and small right (bottom) prism trials for each bat (points); bars indicate the proportion of L-L click-pairs vs. R-R click-pairs averaged over all trials. An asterisk (*) indicates $p < 0.05$ for the Wilcoxon signed-rank test; otherwise, results are not significant (n.s.). Dark bars highlight conditions in which the percentage shifted by >30% when approaching the target. B) (Left) Maximum deviation vs. SD-TD far from (top) and close to (bottom) the target for individual trials across bats. Larger scatter points denote earlier prism trials. Note that early aiming of the beam away from the target is correlated with larger maximum deviations and that earlier trials tend to aim away from the target. (Right) Grey bars show the lowest maximum deviation vs. the lowest SD-TD across bats and trials shaded by prism trial #. Bats learn to aim closer to the target earlier in the trial as the session progresses. Sonar aim remained centered around the apparent visual target later in the trial, not varying significantly over the session. Symbols represent data from different bats.

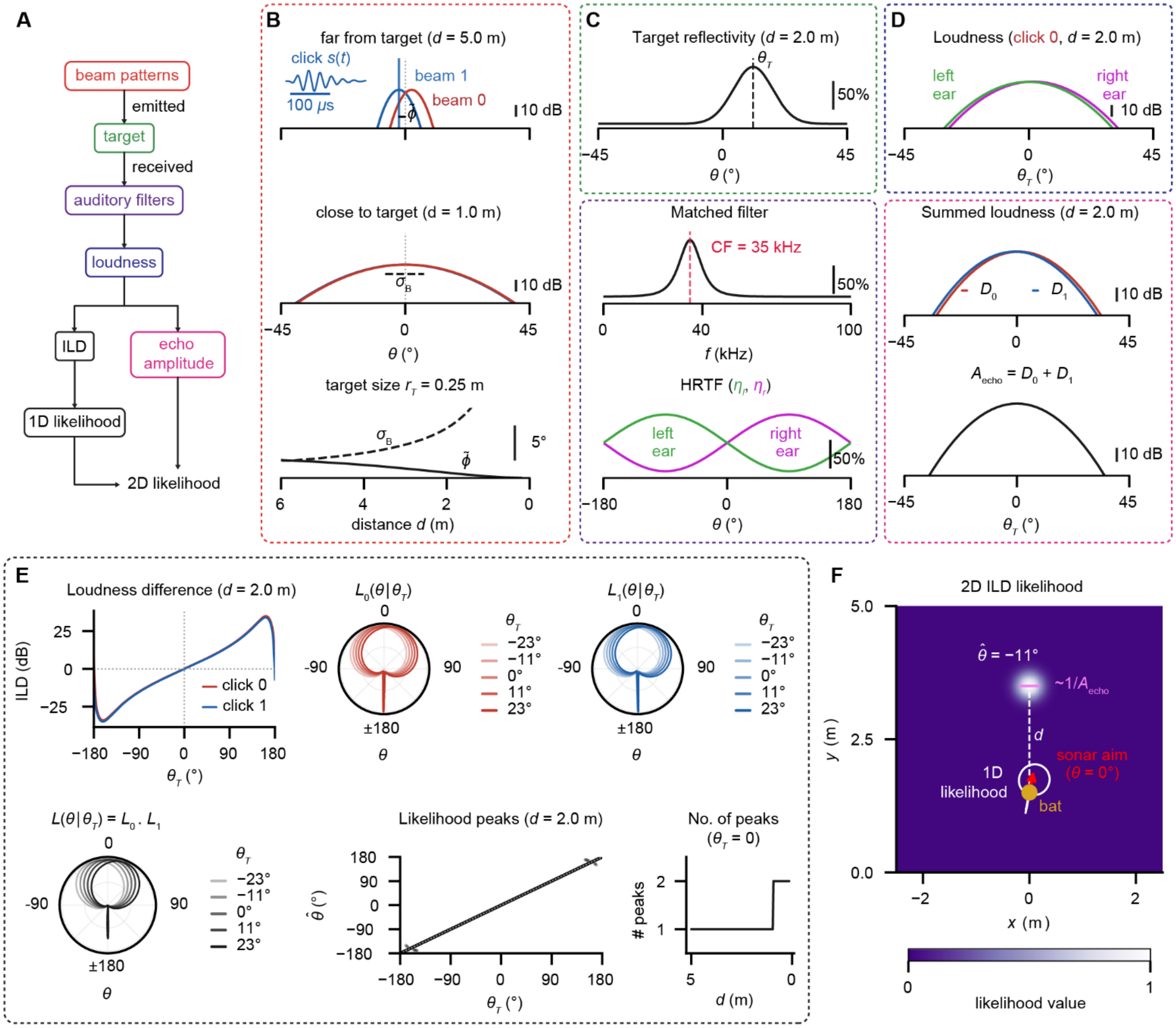

**Fig S7. Model of *Rousettus* echolocation.**

A) Schematic illustrating the sequence of steps from sonar emission to likelihood calculation. B) Beam patterns for a click-pair as a function of angle relative to the sonar aim $\theta$ far from and close to the target. Inset shows the temporal profile of a click modeled as a 40 kHz Gammatone. The beam width $\sigma_B$ and half of the inter-click angle $\phi$ are modulated with the distance to target $d$. C) (Top) Target reflectivity as a function of $\theta$ for a target of size $r_T$ at angle $\theta_T$ relative to sonar aim. (Bottom) Matched auditory filter in frequency and space ("head-related transfer function" or HRTF) as a function of $\theta$. CF is center frequency. D) (Top) Integrated loudness for a single click at the left and right ear. Note how the HRTF creates a difference in the left vs. right loudness as a function of $\theta_T$. (Bottom) Summed loudness for each click and across both clicks ("echo amplitude") as a function of $\theta_T$. E) Inter-auditory loudness difference (ILD) as a function of $\theta_T$. The bat independently computes the likelihood that the target is at an angle $\theta$ relative to the sonar aim given the received ILD (which depends on $\theta_T$) for each click. The fused likelihood $L$ is their product. The maximum-likelihood estimate $\theta$ of $L$ equals $\theta_T$. Note that it can also take an additional spurious value (prominent close to the target) antiparallel to the sonar aim. F) The 2D likelihood function is computed in allocentric coordinates by placing a circular Gaussian with width inversely proportional to the echo amplitude at distance $d$ and angle $\theta$ relative to the bat.

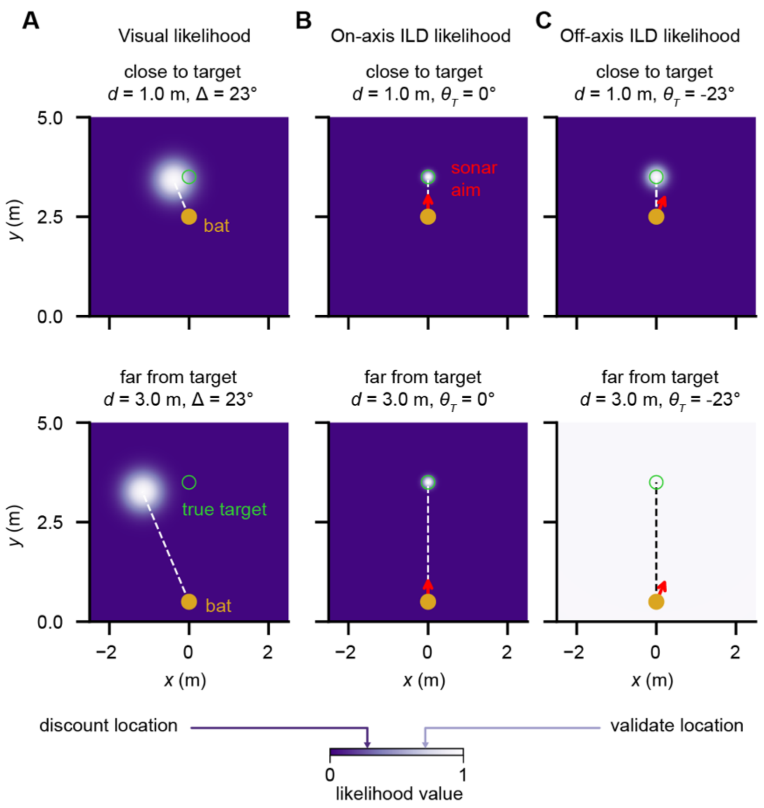

**Fig S8. Model likelihoods depend on distance to target and sonar aim.**
(Top) Likelihood functions close to the target and (Bottom) far from the target. A) Visual likelihoods with the circular Gaussian centered at the true target location rotated counterclockwise by the prism-induced angular shift. B) On-axis ILD likelihoods with the circular Gaussian centered at the true target. C) Off-axis ILD likelihoods. The Gaussian broadens with increasing distance from the true target and increasing angular deviation of the sonar aim from the true target. At sufficiently large distance, off-axis likelihood becomes flat.

**Table S1. Within-subjects Analysis of variance (ANOVA) table comparing hearing sensitivity in each bat (n=4) across sound frequencies with and without ear plugs.**

| | Sum of squares | Mean squares | Degrees of Freedom | F value | P-value |
|---|---|---|---|---|---|
| Condition (plug/no plug) | 17013.6 | 17013.6 | 1, 63 | 191.9173 | **<2.2e-16** |
| frequency | 10397.3 | 1039.7 | 10, 63 | 11.7283 | **4.402e-11** |
| condition x frequency | 1388.2 | 138.8 | 10, 63 | 1.5659 | 0.1379 |

Model: threshold ~ condition * frequency + (1|bat). Degrees of freedom calculated using the Kenward-Rogers method for small samples. Post hoc pairwise testing showed significant differences across plug conditions for all frequencies ($p < 0.05$).

**Movie S1. Model visual and ILD likelihoods along with the posterior belief.**
Visual likelihood (left) instantaneously updates with the apparent location. ILD likelihood (middle) depends on the sonar aim (broad when aim is off axis). Posterior distribution (right) with multiple peaks that determine the sonar aim (in red) and trajectory (in white).

**Movie S2. Bayes risk under a squared-utility error.**
Posterior distribution (left). Peak heights (top right) act as proxies for the probability of visual and auditory hypotheses being true. Loss matrix (middle right) based on the squared-error utility. Bayes risk (bottom right) computed by multiplying the loss matrix with the probabilities. Note that it favors probing the less likely hypothesis.

**Movie S3. Model visual likelihood and posterior belief when acoustic feedback is absent.**
Visual likelihood (left) driven changes in the posterior belief (right) are insufficient for timely course correction.
